# Transition from background selection to associative overdominance promotes diversity in regions of low recombination

**DOI:** 10.1101/748004

**Authors:** Kimberly J. Gilbert, Fanny Pouyet, Laurent Excoffier, Stephan Peischl

**Author notes:** These authors contributed equally (Kimberly J. Gilbert, Fanny Pouyet). Senior authors (Laurent Excoffier, Stephan Peischl).

## Abstract

Linked selection is a major driver of genetic diversity. Selection against deleterious mutations removes linked neutral diversity (background selection, BGS, Charlesworth *et al.* 1993), creating a positive correlation between recombination rates and genetic diversity. Purifying selection against recessive variants, however, can also lead to associative overdominance (AOD, Ohta 1971, Zhao & Charlesworth, 2016), due to an apparent heterozygote advantage at linked neutral loci that opposes the loss of neutral diversity by BGS. Zhao & Charlesworth (2016) identified the conditions when AOD should dominate over BGS in a single-locus model and suggested that the effect of AOD could become stronger if multiple linked deleterious variants co-segregate. We present a model describing how and under which conditions multi-locus dynamics can amplify the effects of AOD. We derive the conditions for a transition from BGS to AOD due to pseudo-overdominance (Ohta & Kimura 1970), i.e. a form of balancing selection that maintains complementary deleterious haplotypes that mask the effect of recessive deleterious mutations. Simulations confirm these findings and show that multi-locus AOD can increase diversity in low recombination regions much more strongly than previously appreciated. While BGS is known to drive genome-wide diversity in humans (Pouyet *et al*. 2018), the observation of a resurgence of genetic diversity in regions of very low recombination is indicative of AOD. We identify 21 such regions in the human genome showing clear signals of multi-locus AOD. Our results demonstrate that AOD may play an important role in the evolution of low recombination regions of many species.

## Results & Discussion

The interaction of recombination with selective and non-selective processes impacts both functional and neutral diversity across the genome, where a positive correlation between genetic diversity and recombination is generally expected (Barton & Etheridge 2004, Charlesworth *et al.* 1993). Yet, Zhao & Charlesworth (2016) have recently identified a possible transition from BGS to AOD at sites linked to a locus with recessive deleterious mutations, leading to a switch from reducing to increasing diversity, which was matched by empirical analysis in *Drosophila* (Becher *et al.* 2019). However, the prevalence of AOD and its impact on genomic diversity in natural populations is unknown, especially in a genomic context where multiple linked recessive loci co-segregate in regions of low recombination. Here, we investigate the evolutionary dynamics of multi-locus AOD through a combination of simulation and mathematical modelling that allows us to identify empirical signatures of AOD in humans.

### Simulations of linked selection

Evidence of background selection (BGS) reducing diversity in regions of low recombination has been observed across many taxa, including *Drosophila* (Begun & Aquadro 1992, Aquadro et al. 1994, Kreitman & Wayne 1994, Shapiro et al. 2006, Kulathinal et al. 2008), humans (Hellman et al. 2005), white-throated sparrows (Huynh et al. 2010), tomatoes (Stephan & Langley 1998), and maize (Tenaillon et al. 2001, 2002). We performed genome-scale simulations to understand how selection (*Ns*), dominance of deleterious alleles (*h*), and recombination rate interact to shape patterns of nucleotide diversity (π) and derived allele frequency (DAFi, Pouyet *et al.* 2018) at neutral sites. For co-dominant mutations, our simulations fully agree with the classical BGS result that linked neutral diversity decreases with decreasing recombination (Figs. 1A & 1B; see e.g. eqn. 8 in Hudson & Kaplan 1995). In agreement with theoretical expectations (e.g. Nordborg *et al.* 1996), BGS has minimal effects on neutral diversity around nearly neutral or strongly deleterious variants (Figs. 1A & 1B), since selection is either so weak (*s* = -0.00001) that it has little impact on fitness, or so strong (*s* = -0.1) that mutations are rapidly purged before diversity can be reduced (Supplemental Figure S1).

**Figure 1.**
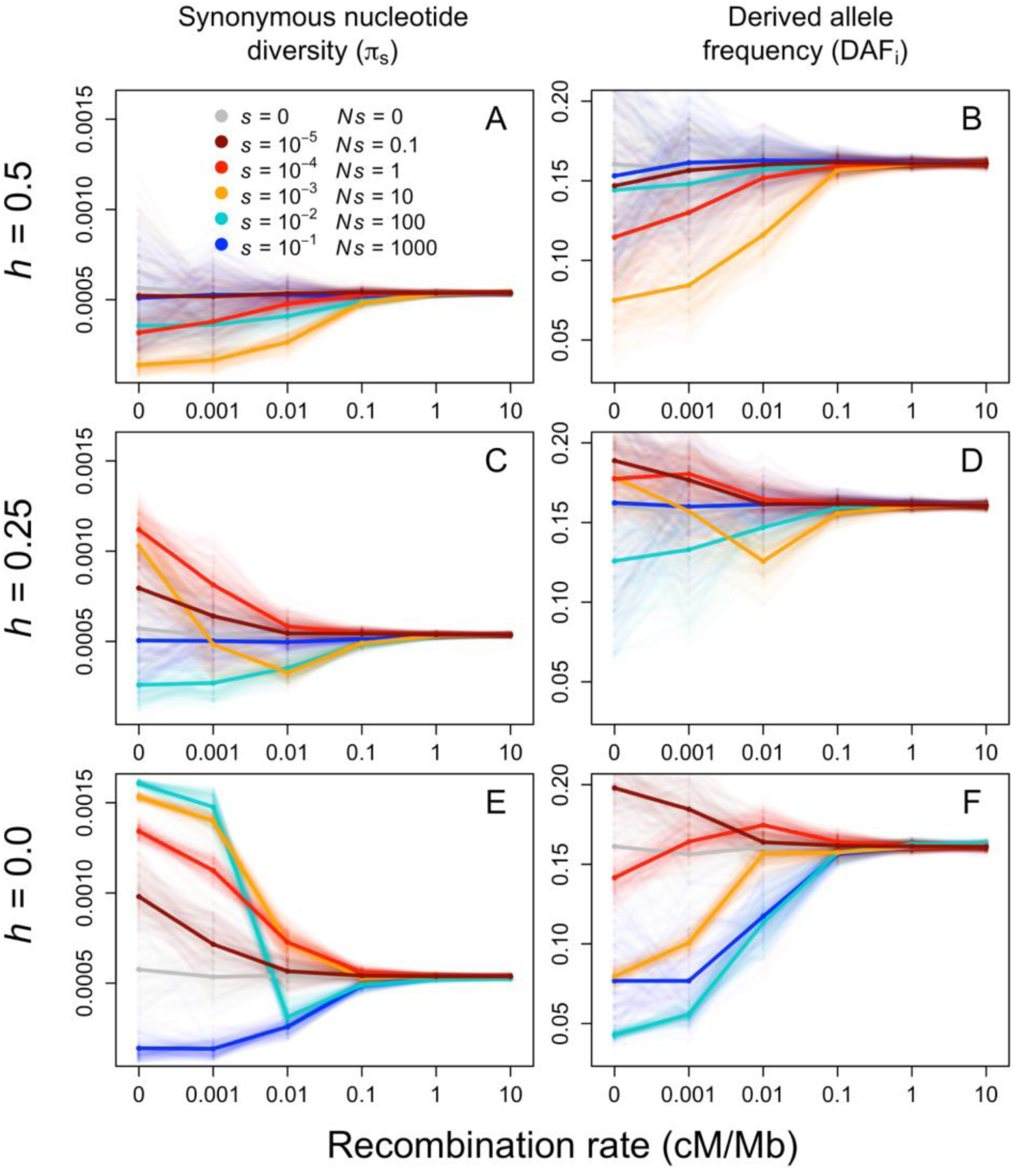
Genomic diversity at neutral sites as measured by nucleotide diversity (π) and derived allele frequency (DAF_i_) from simulations across cases of selection coefficients (s), dominance of deleterious alleles (h), and recombination rate. Means across all replicates are shown in thick lines while all 100 replicates are transparent lines in the background.

In the presence of partially (*h* = 0.25) and fully (*h* = 0) recessive deleterious alleles, neutral nucleotide diversity (π) increases at lower recombination rates (Figures 1C & 1E), except when selection is very strong (*Ns* = 1000). In some cases, we observe that nucleotide diversity decreases and then rebounds as recombination decreases (e.g. *Ns* = 10, orange lines in Figure 1C), potentially reflecting a transition from BGS to AOD. The purging of homozygous genotypes in regions of low recombination is expected to only slightly increase diversity (on the order of a few percent) and be limited to small selection coefficients (*Ns* ≲ = 1, Zhao & Charlesworth 2016, but see Becher *et al.* 2019). In our simulations, however, we often see more than a two-fold increase in diversity, even for selection coefficients where we would expect a reduction of diversity due to BGS (e.g., for *Ns* = 100 in Figure 1E). Our results thus extend the finding of Zhao & Charlesworth (2016) that two tightly linked loci under selection show stronger AOD than predicted by single-locus theory, suggestive of a synergistic effect between linked deleterious sites.

We gain further insight by contrasting π with the derived allele frequency per individual (DAFi, see STAR Methods). Nucleotide diversity captures the amount of heterozygosity in the population whereas DAFi measures allele frequencies at all polymorphic loci, and thus the presence of rare heterozygotes affects these two quantities in contrasting ways. We find that qualitatively different patterns of diversity are observed across π and derived allele frequencies for various selection intensities (Figure 1C-F). For example, in Figure 1E, the greatest π and thus greatest impact of AOD is obtained when *Ns* = 100, *h* = 0, and no recombination, yet this is equivalently the lowest respective DAFi measure across simulated cases. Contrasting DAFi and π thus suggests that AOD can lead to maintenance of high levels of diversity via many polymorphic sites at low frequency (high π, low DAFi, e.g. *Ns* = 100 in Figs 1E & 1F), or via few sites with high derived frequencies (high π, high DAFi, e.g. *Ns* = 0.1 in Figs 1E & 1F).

In agreement with this explanation the SFS show peaks of intermediate frequency variants (Fig. 2), with most pronounced peaks for intermediate selection coefficients. As selection coefficients increase, peaks shift to lower classes of the SFS and become more pronounced (Figure 2C-D). Clustering analyses suggest that the increased heterozygosity is caused by maintenance of complementary deleterious haplotypes in the population (Supplemental Figure S2A-C). Thus, higher selection coefficients imply a larger number of rare complementary haplotypes across loci and therefore higher π but lower DAFi values. For smaller selection coefficients, both π and DAFi are high because complementary haplotypes are maintained in the population at higher frequencies (Figures 1E-F, 2A-B). Meanwhile under our strongest selection case (*Ns* = 1000), AOD is not observed, as we explain in the following model.

**Figure 2.**
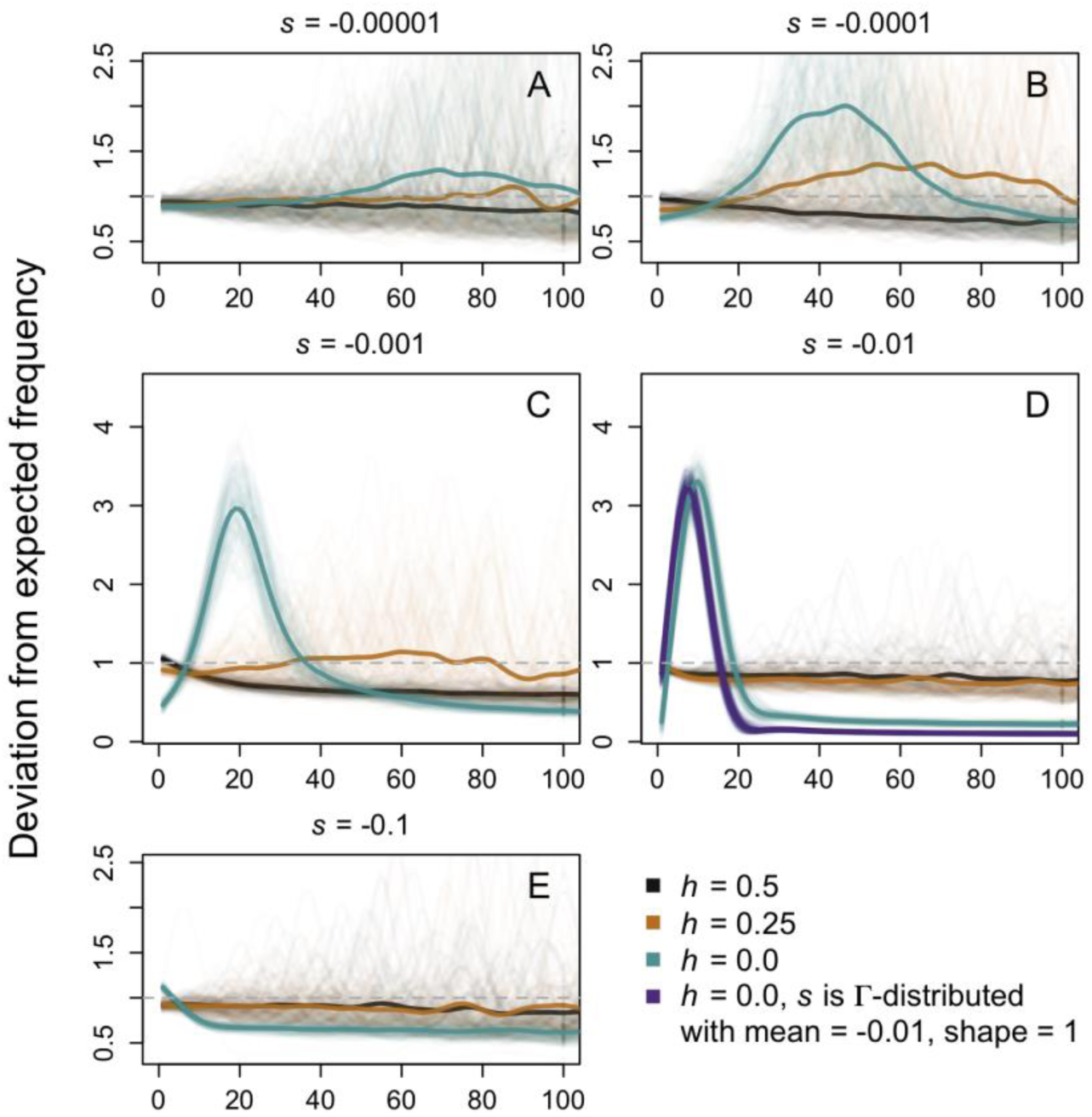
Site frequency spectra for neutral sites calculated relative to the expected SFS across the whole genome (all recombination rates). Data is smoothed with a spline function, average SFS across all SLiM simulations are shown in thick solid lines, and individual replicates are translucent lines in background. Only the first 100 frequency classes of the SFS are shown (from n=1-100) because classes from n=101-200 add no information and lines are flat.

### Modelling the transition from BGS to AOD

To better understand the action of linked selection on neural variants, we developed an analytical model of the multi-locus dynamics of mutation, recombination, and selection acting on fully recessive variants. Our goal is to find the conditions for a transition from BGS to selection maintaining complementary haplotypes. This complementation arises as selection favors multi-locus heterozygotes over individuals homozygous for recessive variants (pseudo-overdominance, Ohta & Kimura 1970), creating multi-locus AOD at nearby neutral sites. Pseudo-overdominance occurs only when the wild-type haplotype without any deleterious mutations is lost from the population, that is, when every individual carries at least one deleterious allele and heterozygotes are fitter than any of the homozygotes. Frequent recombination would re-create the wild-type haplotype, and we thus expect a transition from BGS to pseudo-overdominance and strong AOD only when recombination is sufficiently weak.

We consider a region with map length *r* that harbors *n* equidistant sites at which deleterious mutations occur at rate *u* per site. In Supplemental Item 1, we show that in an (effectively infinitely) large population the frequency of the wild-type class at mutation-selection balance can be approximated by 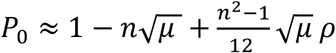, where 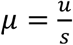 and 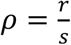 are the scaled strengths of mutation and recombination, respectively. We use this result to predict the conditions for loss of the wild-type haplotype (that is, when *P*_0_ = 0) and hence a transition from purifying selection to pseudo-overdominance, which fits well to two-locus simulations (Figure 3A). Figure 3B shows that increasing the number of deleterious loci while keeping the per-locus mutation rate constant increases the probability of losing of the wild-type haplotype if the recombination rate is very low, simply because the total deleterious mutation rate (increases. With higher recombination, loss of the wild-type haplotype is most likely for an intermediate number of deleterious sites (Figure 3B). The reason is that with multiple segregating loci recombination is more likely to produce the wild-type haplotype and is thus more efficient at preventing a transition to AOD.

**Figure 3.**
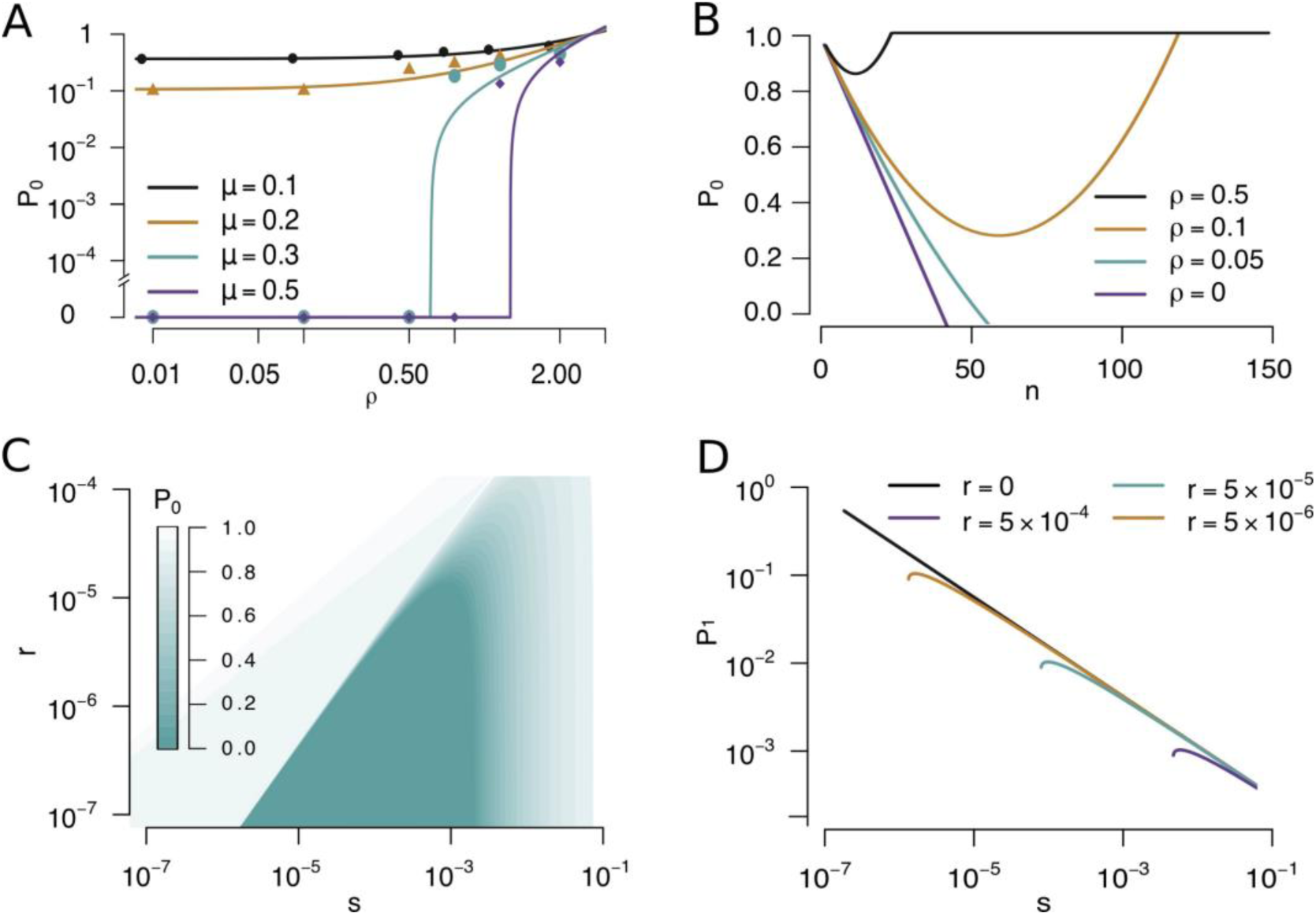
Frequency of the wild-type haplotype at mutation-selection balance. (A) Individual-based simulations for a two-locus model with parameters s = 0.001, h = 0, N = 10 000 diploid individuals, and μ and ρ as specified in the figure. Solid lines show analytical theory and dots the results from simulations. (B) Expected frequency of the wild-type haplotype as a function of the of deleterious loci n. Parameters values are s = 10^-4^, h = 0, and u = 1.4510^-8^. (C) Combinations of r and s for which a transition from BGS to AOD is predicted. Parameters are u = 1.45*10^-8^, n = 250. The parameter r measures the recombination rate between two consecutive deleterious sites. (D) Predicted equilibrium frequency of each complementary haplotype (P_1_) after the transition to pseudo-overdominance. Note that this is also the expected per-locus allele frequency at selected loci because every haplotype carries a unique deleterious mutation. The colored lines show the haplotype frequency if there is recombination. For small values of s and positive recombination rates, the lines end because if recombination is sufficiently strong relative to selection there is no transition to AOD.

Loss of the wild-type haplotype is most likely for intermediate selection coefficients (Figure 3C), explaining why we observe the strongest increase in diversity for intermediate selection coefficients (Figure 1E). When mutations are only slightly deleterious, recombination is too strong relative to selection and the wild-type haplotype is unlikely to be lost. Alternatively, if selection is very strong relative to the mutation rate, haplotypes carrying a deleterious allele will be purged quickly and loss of the wild-type haplotype is unlikely, explaining our observation of strictly decreasing diversity under the strongest selection coefficient simulated (dark blue lines, Fig. 1E-F). Once the wild-type haplotype is lost, complementary haplotypes will be maintained by pseudo-overdominance (see Supplemental Figure S3), and our model predicts that the equilibrium frequency of these haplotypes decreases with increasing selection coefficients. Our model predictions closely match our simulations: nucleotide diversity will generally be high under multi-locus AOD due to pseudo-overdominance that increases heterozygosity. DAFi will, however, only be large if selection coefficients are small enough because the frequency of complementary haplotypes, and hence the per-locus derived allele frequency, decreases with increasing selection intensity (Figure 3D). Our analytical results also explain why a strong increase in heterozygosity is not necessarily linked to an increase in DAFi if selection is strong (e.g., *s* = −0.0 in Figures 1E and 1F).

### Empirical Inference of AOD

Our theoretical results indicate that multi-locus AOD generally increases π in regions of low recombination. Based on this result, we scan 100 human genomes from ten 1000G populations to find regions subject to multi-locus AOD (see STAR Methods for details). Overall, we identified 21 regions (33 windows, some partially overlapping) where the difference in π between low and medium recombination rates exceeds two standard deviations around the chromosomal mean across all ten populations (Supplemental Item 2A, Supplemental Table S1). Among these 21 regions, 18 overlap with those identified by Bitarello *et al.* (2018) and hypothesized to be under long-term balancing selection based on a statistic classifying frequency classes within the SFS. However, disentangling between ‘classical’ balancing selection (single-locus overdominance), and pseudo-overdominance is difficult as both processes leave similar signatures of genomic diversity in low recombination regions. For instance, our scan detects the Major Histocompatibility Complex (MHC) as a candidate region for AOD, yet it is a region known to be under long-term balancing selection and not subject to pseudo-overdominance (Meyer & Thomson, 2001). Contrastingly, the region we identify with the highest difference in π between low and medium recombination (Supplemental Table S1) is located on Chromosome 9:31,000,001-33,500,000 (Figure 4A) and is a promising candidate for multi-locus AOD. This region contains the gene SPINK4 (Serine peptidase inhibitor, Kazal type 4; Supplemental Table S2 shows results from functional analysis of this outlier region), which has recently been identified in a scan for sites under balancing selection (Bitarello *et al.* 2018). SPINK4 belongs to the Serpin Superfamily of proteins which inhibit target proteases via a conformational change that disrupts the active site (Whisstock & Bottomley 2006). Serpins are hypothesized to be under purifying selection since a mutation could lead to protein aggregation and a disease known as serpinopathy (Carrell & Lomas, 1997). The candidate region contains other functionally important genes potentially under purifying selection (Supplemental Table S2) such as Y RNA that are involved in DNA replication (Christov *et al.* 2006) and in the regulatory activity of Ro60, a quality control protein which targets misfolded small RNA (Chen & Wolin, 2004). The presence of several functionally important genes that could contain many slightly deleterious alleles suggests that higher diversity and the signature of AOD in this region may be driven by pseudo-overdominance rather than overdominance. AOD is thus a plausible explanation for the odd observation that many variants found in scans for balancing selection are associated with severe consequences in protein-coding genes (Key *et al.* 2014).

**Figure 4.**
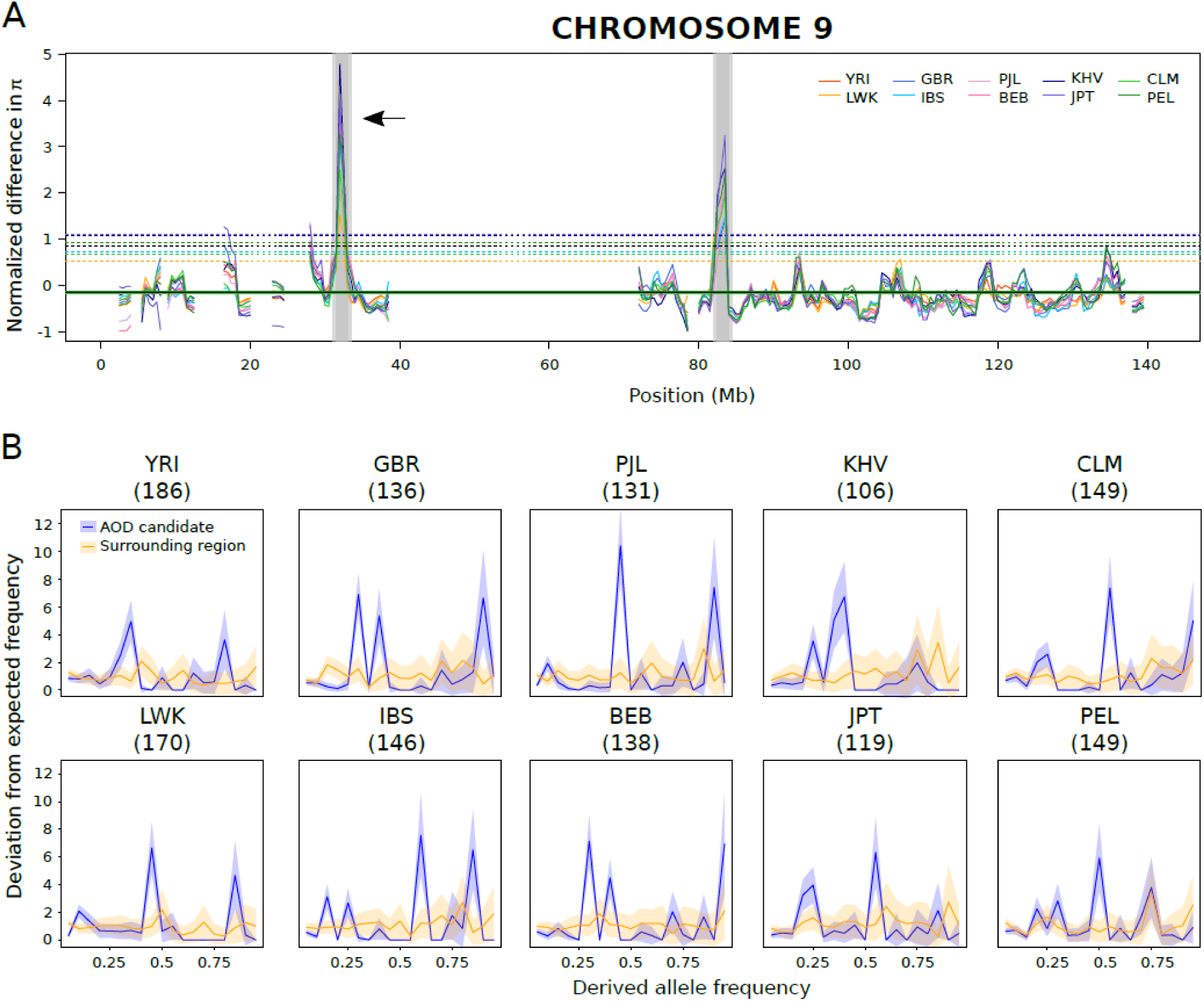
Genome scan of the normalized difference in π between low and medium recombination on Chromosome 9 for ten human populations. (A) The shaded grey area represents candidate windows for AOD i.e. the normalized difference is higher than two standard deviations (dashed lines) from the mean for each population (see STAR Methods for details). The arrow indicates the outlier with the highest average normalized difference across 10 populations, which is focused on in the text and panel B. (B) Normalized SFS across the ten populations of low recombination (LR) SNPs in the candidate region (blue) versus LR SNPs in the surrounding 5Mb region (orange). Number of sampled variants are indicated in parentheses. Confidence intervals (shaded areas) are estimated from 100 bootstrap replicates.

Additional support for multi-locus AOD comes from the fact that the SFS show strong peaks at intermediate frequencies for all populations in the candidate regions compared to the surrounding ones (Figure 4B, Supplemental Item 2B). While such peaks could be indicative of a mapping problem, we excluded such potentially problematic genomic regions by using the most stringent genomic quality masks (see STAR Methods). Moreover, we find noticeable clustering in the derived status of many loci across sampled individuals, suggestive of complementary haplotypes (Supplemental Item 2C), particularly in non-African populations.

## Conclusions

Our results show that multi-locus AOD resulting from pseudo-overdominance of deleterious variants has the potential to impact neutral genomic diversity in regions of low recombination. Importantly, the resulting increase in diversity is much stronger than previously identified from one- or two-locus models and can lead to a form of selection maintaining complementary heterozygous haplotypes. The conditions under which we expect a transition from BGS to multi-locus AOD depend on the dominance of deleterious alleles, the strength of selection, and the recombination rate. The rate of influx of new deleterious mutations also plays a critical role, since there is a balance between strength of selection, the rate of recombination and the speed at which these variants are removed (Figure 3A, Supplemental Item 2C). These interactions thus determine whether deleterious variants are favored in the heterozygous state or if selection is able to efficiently remove them before they accumulate in heterozygotes across the genome, creating a resurgence of genetic diversity in regions of low recombination.

Intriguingly, multi-locus AOD within the genome is differentially detected by π and DAFi depending on the strength of linked selection and the degree of dominance. Contrasting π and DAFi could thus open avenues for statistical inference of the joint distribution of selection and dominance coefficients. Empirical investigation into the human genome finds that among the 21 regions likely impacted by multi-locus AOD, 19 have a higher DAFi at low versus intermediate recombination rates (Supplemental Table S1). Our theoretical results would thus indicate that most selected variants in these regions are slightly deleterious (e.g. Figure 1D & 1F, *Ns* = 0.1). Variable demography and genetic drift may impact the dynamics of AOD in the genome. For instance, the transition from BGS to AOD through loss of the wild-type haplotype and increase in frequency of recessive deleterious variants could become more likely due to pulses of increased genetic drift, such as bottlenecks and range expansions. Along these lines, all 21 candidate regions show a clear excess of variants at intermediate frequencies, yet different populations exhibit variability in the SFS (Figure 4B, Supplemental Item 2B). Generally we find stronger signals of multi-locus AOD in non-African populations, both in the SFS with higher frequency peaks as well as in heat maps of haplotype structure showing larger clusters of derived alleles (Figure 4B, Supplemental Items 2B & 2C).

Understanding the causes, consequences, and conditions leading to BGS and AOD is a major step in understanding the drivers of diversity across the genome. Our results inform how selection against deleterious variants can impact the evolution of linked neutral variation and lead to unexpected patterns of diversity in regions of low recombination. Inference methods making use of SFS data may thus be biased if regions of low recombination are significant contributors to measured diversity, emphasizing the need for better understanding of the impacts of linked selection in the genome. For instance, multi-locus AOD may lead to misidentification in scans for balancing selection, interfere with the detection of introgressed or inverted regions of the genome where large regions of heterozygosity are expected, or mimic signatures of local adaptation detected by *FST* outlier scans. This study opens promising avenues for further empirical and theoretical research into the interaction of recombination with drivers of genomic diversity, including the efficiency of selection against introgressed variation (Kelley & Nielsen 2016), evolutionary dynamics of mutation load and inbreeding depression, and evolution of recombination modifiers (Berdan *et al*. 2019).

## Acknowledgements

We would like to thank Brian Charlesworth, Vitor Sousa, and Claudia Bank for useful insights, discussion, and feedback on the topics of this paper. FP and KJG were funded by a Swiss NSF grant No 310030B-166605 to LE.

## Author Contributions

KJG, FP, LE, and SP conceived the study. KJG performed all simulation analyses, SP developed all analytic theory, and FP performed all human analyses. KJG, FP, and SP wrote the first draft of the paper, and all authors made significant contributions to the final draft.

## STAR Methods

### Key Resources Table

| REAGENT or RESOURCE | SOURCE | IDENTIFIER |
| --- | --- | --- |
| Deposited Data |  |  |
| Human variants | The 1000G project Consortium, 2015 | http://hgdownload.cse.ucsc.edu/gbdb/hg19/1000Genomes/phase3/ |
| Human accessible genome masks | The 1000G Project Consortium, 2015 | ftp://ftp.1000genomes.ebi.ac.uk/vol1/ftp/release/20130502/supporting/accessible_genome_masks/ |
| Recombination map | The International HapMap Consortium, 2007 | ftp://ftp.1000genomes.ebi.ac.uk/vol1/ftp/technical/working/20130507_omni_recombination_rates/ |
| Derived alleles frequencies per individual per population | Pouyet et al., 2018 | https://datadryad.org/resource/doi:10.5061/dryad.t76fk80.2/16.2 |
| Software and Algorithms |  |  |
| SLiM v3.2 | Haller & Messer 2019 | https://messerlab.org/slim/ |
| Mathematica | Wolfram | http://www.wolfram.com/mathematica/ |
| bedtools | Quilan, 2014 | https://github.com/arq5x/bedtools2/releases/download/v2.28.0/bedtools-2.28.0.tar.gz |
| vcftools | Danecek *et al.*, 2011 | https://sourceforge.net/projects/vcftools/files/ |
| BioMart | Smedley *et al.*, 2015 | http://grch37.ensembl.org/biomart/martview/1a32c1634d968649752622cb3c48081d |
| Other |  |  |
| Simulation analysis files (R scripts) | This paper | To be stored on Github |
| Mathematica code/scripts | This paper | To be stored on Github |
| Human genome analysis scripts | This paper | https://github.com/FannyPouyet/mAOD |

### Contact for Reagent and Resource Sharing

Further information and requests for resources should be directed to and will be fulfilled by the Lead Contact, Kimberly J. Gilbert.

### Method Details

#### Simulations

We conduct forward-time, individual-based simulations to clarify the processes of BGS and AOD occurring in the genome, using SLiM v3.2 (Haller & Messer 2019). We model a census population size of 10,000 individuals in one panmictic, constant-sized population, and sample 100 individuals from this population at generation 6*N*. The per-bp mutation rate in all simulations is 1.45e^-8^, matched to latest estimates from humans (Scally 2016), and we prevented mutations from stacking (see SLiM manual). Only mutations unconditionally deleterious to fitness or neutral are considered, *i.e.* we do not include the impact of beneficial mutations or potential selective sweeps that would be associated with such variants.

Individuals consist of six chromosomes of 7.5Mb each, making each individual’s genome a total size of 45Mb. Each chromosome is subject to only one recombination rate (0, 0.001, 0.01, 0.1, 1, and 10 cM/Mb), and each replicate simulation only exhibits deleterious mutations of one constant, fixed selection coefficient *s* (−0.00001, -0.0001, -0.001, -0.01, -0.1). One set of simulations modelled a gamma distribution for deleterious fitness effects with mean *s* = -0.01 and shape parameter 1, i.e. an exponential distribution. Mutations are only either neutral or deleterious, and we varied the proportion of deleterious mutations to be either 2% or 20% (Supplemental Figure S4) of the total mutations occurring at any given point in time. Across replicates we also varied the dominance of deleterious mutations (*h* = 0.5, 0.25, 0; additive, partially recessive, and fully recessive, respectively).

#### Mathematical model

We consider *n* biallelic loci and assume these loci are equidistant from each other with respect to recombination distance. We denote the recombination rate between the first and last locus as *r* Thus, the recombination distance between two adjacent loci is approximately *r*/(*n* − 1). Derived alleles are recessive and deleterious with fitness effect *s* in the homozygous state. We assume that fitness is multiplicative across loci, *i.e.*, there are no epistatic effects across loci. Mutations occur at each locus with rate per generation and individual, and *U*_*d*_ = *n u* is the total genome-wide deleterious mutation rate.

#### Human Analyses

We use a genotype table containing 100 individuals from ten 1000G populations (The 1000G Project Consortium, 2015) from Pouyet *et al*. (2018). Population labels are according to The 1000G Project Consortium nomenclature: YRI = Yoruba, LWK = Luhya, GBR = British, IBS = Spanish, BEB = Bengali, PJL = Punjabi, KHV = Vietnamese, JPT = Japanese, CLM = Colombian, and PEL = Peruvian (names of individuals are available with the archived analyses on GitHub). We use bedtools (Quilan 2014) to focus on genomic regions with very low negative and positive false discovery rates from next generation sequencing methods by using the 1000G strictMask map. This map keeps genomic regions where the depth of coverage (summed across all 1000G samples) is within 50% of the average, where no more than 0.1% of reads have a mapping quality of zero and where the mapping quality is > 56 (The 1000G Project Consortium, 2015). This generates a total of 13,385,820 SNPs. We use the LD-based Yoruba-specific recombination map (The International HapMap Consortium, 2007) to define sites in regions of low recombination (LR) with 0 < RR <= 0.05 cM/Mb and medium recombination (MR) with 1 <= RR < 1.5 cM/Mb.

### Quantification and Statistical Analysis

#### Summary statistics

To understand the evolutionary dynamics across our combinations of simulated parameters, we compare several summary statistics from polymorphic sites in the simulated data. The derived allele frequency (DAFi) in the population sample is calculated according to Pouyet *et al.* (2018), across only neutral, polymorphic sites. We generate the unfolded site frequency spectrum (SFS) from the sample of 100 diploid individuals, across the whole genome (*i.e.* including sites subject to all recombination levels). This SFS is normalized against the expected SFS for an ideal population (Lapierre *et al.* 2017) by dividing each class by the sum of all 200 classes, creating an SFS where deviation from 1 indicates an enrichment or deficit of variants in that SFS class.

#### Identification of AOD in Humans

We perform a window-based approach with window sizes of 2Mb and a step increase of 500kb to scan all chromosomes of the human genome per population. We kept only windows with at least 50 SNPs for both LR and MR and compute the average derived allele frequency (DAFi) across the 10 populations for LR and MR. We use vcftools (Danecek *et al.* 2011) to estimate nucleotide diversity (π) per site and average it per window using a custom script for LR and MR. For each chromosome and each window *i*, we estimate the difference in nucleotide diversity between LR and MR per population:

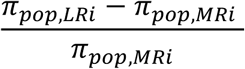

Windows are averaged across chromosome per population, and outliers are identified as those exceeding two standard deviations around this mean in all ten populations. Finally, we used biomart (Smedley *et al.* 2015) to identify the gene functions present in these regions, as an analysis of AOD due to balancing selection versus pseudo-overdominance.

#### Data and Code Availability

No new data was generated in this study. Input code and scripts for theoretical and simulation results are archived as described in the Key Resources Table, in publicly accessible repositories.

