## Supplemental Material for "Transition from background selection to associative overdominance promotes diversity in regions of low recombination"

### Supplemental Information

**Supplemental Table S1.** Summary of the 21 genome-wide regions identified as AOD candidates, with the coordinates, number of SNPs in low and medium recombination regions (LR and MR respectively), and the average for the 10 populations of the normalized difference in π, $DAF_i$ for low recombination ($0 < RR \leq 0.05$ cM/Mb), and $DAF_i$ for medium recombination ($1 \leq RR < 1.5$ cM/Mb).

| Chromosome | START | END | Num. variants with LR | Num. variants with MR | $\overline{\frac{\pi_{LR} - \pi_{MR}}{\pi_{MR}}}$ | $\overline{DAFi_{LR}}$ | $\overline{DAFi_{MR}}$ |
|---|---|---|---|---|---|---|---|
| 1 | 64500001 | 66500000 | 777 | 355 | 0.638673 | 0.165972 | 0.154859 |
| 1 | 2.36E+08 | 2.38E+08 | 353 | 1062 | 1.438257 | 0.232238 | 0.170621 |
| 2 | 2.43E+08 | 2.45E+08 | 143 | 68 | 0.732335 | 0.152762 | 0.215882 |
| 3 | 61500001 | 63500000 | 106 | 1104 | 1.715917 | 0.20033 | 0.148007 |
| 3 | 1.89E+08 | 1.91E+08 | 115 | 1456 | 1.075936 | 0.22313 | 0.153644 |
| 4 | 23500001 | 26500000 | 63 | 1588 | 1.365892 | 0.254365 | 0.129005 |
| 5 | 84000001 | 8.60E+07 | 1100 | 245 | 0.625868 | 0.168691 | 0.148204 |
| 6 | 19000001 | 2.20E+07 | 252 | 783 | 1.182227 | 0.189365 | 0.171616 |
| 6 | 32000001 | 3.40E+07 | 2831 | 647 | 0.959907 | 0.231676 | 0.176917 |
| 7 | 1000001 | 3.00E+06 | 115 | 370 | 1.600572 | 0.142522 | 0.12577 |
| 8 | 1.23E+08 | 1.25E+08 | 173 | 358 | 2.440507 | 0.262197 | 0.131103 |
| 9 | 31000001 | 33500000 | 261 | 721 | 3.259377 | 0.196877 | 0.176865 |
| 9 | 82000001 | 84500000 | 101 | 277 | 1.593002 | 0.228218 | 0.145307 |
| 10 | 46500001 | 49500000 | 81 | 174 | 2.043011 | 0.232778 | 0.150862 |
| 10 | 1.12E+08 | 1.14E+08 | 248 | 703 | 1.789422 | 0.232661 | 0.145064 |
| 11 | 96500001 | 99500000 | 219 | 1000 | 3.14817 | 0.226849 | 0.16427 |
| 12 | 1.3E+08 | 1.32E+08 | 94 | 2007 | 1.123755 | 0.155851 | 0.166355 |
| 13 | 63000001 | 6.50E+07 | 1905 | 160 | 0.789565 | 0.164215 | 0.095563 |
| 16 | 1 | 3500000 | 128 | 1310 | 2.162051 | 0.260977 | 0.145004 |
| 18 | 40500001 | 42500000 | 193 | 254 | 1.235513 | 0.19057 | 0.123661 |
| 21 | 41000001 | 4.30E+07 | 100 | 1218 | 1.685968 | 0.29935 | 0.189906 |

**Supplemental Table S2.** Genic analysis of the outlier window on chromosome 9 (31,000,001 to 33,500,000 bp). This window is a candidate for AOD due to pseudo-overdominance with 52 genes identified with Biomart – Ensembl (see STAR Methods). lincRNA corresponds to long intergenic non coding RNA and MiRNA to small-non coding RNA which are affecting genes expression via silencing or post-transcriptional regulation.

| Gene stable ID | Gene name | Gene description |
|---|---|---|
| ENSG00000223421 | RP11-572H4.1 | Pseudogene |
| ENSG00000232203 | SLC25A6P2 | solute carrier family 25 (mitochondrial carrier; adenine nucleotide translocator), member 6 pseudogene 2 [Source:HGNC Symbol;Acc:43850] |
| ENSG00000260720 | RP11-271O3.1 | LincRNA |
| ENSG00000226998 | RP11-291J9.1 | Pseudogene |
| ENSG00000229080 | RP11-291J9.2 | Pseudogene |
| ENSG00000235148 | HMGB3P23 | high mobility group box 3 pseudogene 23 [Source:HGNC Symbol;Acc:39315] |
| ENSG00000252313 | RNA5SP281 | RNA, 5S ribosomal pseudogene 281 [Source:HGNC Symbol;Acc:43181] |
| ENSG00000225347 | SLC25A5P8 | solute carrier family 25 (mitochondrial carrier; adenine nucleotide translocator), member 5 pseudogene 8 [Source:HGNC Symbol;Acc:35469] |
| ENSG00000122729 | ACO1 | aconitase 1, soluble [Source:HGNC Symbol;Acc:117] |
| ENSG00000107201 | DDX58 | DEAD (Asp-Glu-Ala-Asp) box polypeptide 58 [Source:HGNC Symbol;Acc:19102] |
| ENSG00000197579 | TOPORS | topoisomerase I binding, arginine/serine-rich, E3 ubiquitin protein ligase [Source:HGNC Symbol;Acc:21653] |
| ENSG00000203543 | AL353671.2 | Pseudogene |
| ENSG00000203542 | AL353671.1 | Pseudogene |
| ENSG00000235453 | TOPORS-AS1 | TOPORS antisense RNA 1 [Source:HGNC Symbol;Acc:31420] |
| ENSG00000165264 | NDUFB6 | NADH dehydrogenase (ubiquinone) 1 beta subcomplex, 6, 17kDa [Source:HGNC Symbol;Acc:7701] |
| ENSG00000203659 | AL353671.3 | Pseudogene |
| ENSG00000214006 | AL353671.4 | Pseudogene |
| ENSG00000232303 | RP11-205M20.7 | Pseudogene |
| ENSG00000241043 | GVQW1 | GVQW motif containing 1 [Source:HGNC Symbol;Acc:31424] |
| ENSG00000122728 | TAF1L | TAF1 RNA polymerase II, TATA box binding protein (TBP)-associated factor, 210kDa-like [Source:HGNC Symbol;Acc:18056] |
| ENSG00000256239 | AL589642.1 | Pseudogene |
| ENSG00000223440 | RP11-555J4.4 | Antisense |
| ENSG00000230516 | RP11-555J4.3 | LincRNA |
| ENSG00000238001 | RP11-555J4.2 | Pseudogene |
| ENSG00000230867 | RP11-462B18.1 | Pseudogene |
| ENSG00000221696 | AL157884.1 | MiRNA |
| ENSG00000188133 | TMEM215 | transmembrane protein 215 [Source:HGNC Symbol;Acc:33816] |
| ENSG00000272113 | Y_RNA | Y RNA [Source:RFAM;Acc:RF00019] |
| ENSG00000236796 | RP11-462B18.3 | Pseudogene |
| ENSG00000231193 | RP11-462B18.2 | LincRNA |
| ENSG00000223807 | RP11-562M8.4 | Pseudogene |
| ENSG00000237437 | ASS1P12 | argininosuccinate synthetase 1 pseudogene 12 [Source:HGNC Symbol;Acc:762] |
| ENSG00000137074 | APTX | aprataxin [Source:HGNC Symbol;Acc:15984] |
| ENSG00000236184 | TCEA1P4 | transcription elongation factor A (SII), 1 pseudogene 4 [Source:HGNC Symbol;Acc:31091] |
| ENSG00000225693 | RP11-54K16.2 | Pseudogene |
| ENSG00000086061 | DNAJA1 | DnaJ (Hsp40) homolog, subfamily A, member 1 [Source:HGNC Symbol;Acc:5229] |
| ENSG00000122692 | SMU1 | smu-1 suppressor of mec-8 and unc-52 homolog (C. elegans) [Source:HGNC Symbol;Acc:18247] |
| ENSG00000222169 | AL162590.1 | MiRNA |
| ENSG00000086062 | B4GALT1 | UDP-Gal:betaGlcNAc beta 1,4- galactosyltransferase, polypeptide 1 [Source:HGNC Symbol;Acc:924] |

| | | |
|---|---|---|
| ENSG00000233554 | RP11-326F20.5 | Antisense |
| ENSG00000252224 | RNU4ATAC15P | RNA, U4atac small nuclear 15, pseudogene [Source:HGNC Symbol;Acc:46901] |
| ENSG00000122711 | SPINK4 | serine peptidase inhibitor, Kazal type 4 [Source:HGNC Symbol;Acc:16646] |
| ENSG00000107262 | BAG1 | BCL2-associated athanogene [Source:HGNC Symbol;Acc:937] |
| ENSG00000086065 | CHMP5 | charged multivesicular body protein 5 [Source:HGNC Symbol;Acc:26942] |
| ENSG00000086102 | NFX1 | nuclear transcription factor, X-box binding 1 [Source:HGNC Symbol;Acc:7803] |
| ENSG00000200544 | Y_RNA | Y RNA [Source:RFAM;Acc:RF00019] |
| ENSG00000165269 | AQP7 | aquaporin 7 [Source:HGNC Symbol;Acc:640] |
| ENSG00000223678 | RP11-311H10.4 | Antisense |
| ENSG00000165272 | AQP3 | aquaporin 3 (Gill blood group) [Source:HGNC Symbol;Acc:636] |
| ENSG00000270542 | RP11-311H10.7 | Pseudogene |
| ENSG00000165271 | NOL6 | nucleolar protein 6 (RNA-associated) [Source:HGNC Symbol;Acc:19910] |

**Figure S1. Age and count of segregating variants.**

The mean generation that deleterious mutations arose in the simulations (per replicate and averaged for the thick solid line, for neutral variants in the case of *s* = 0) and the average count of segregating neutral sites in the population. The simulations are forward-time from 0 to 60,000 generations, making mutations arising at generation 60,000 the youngest mutations in the population.

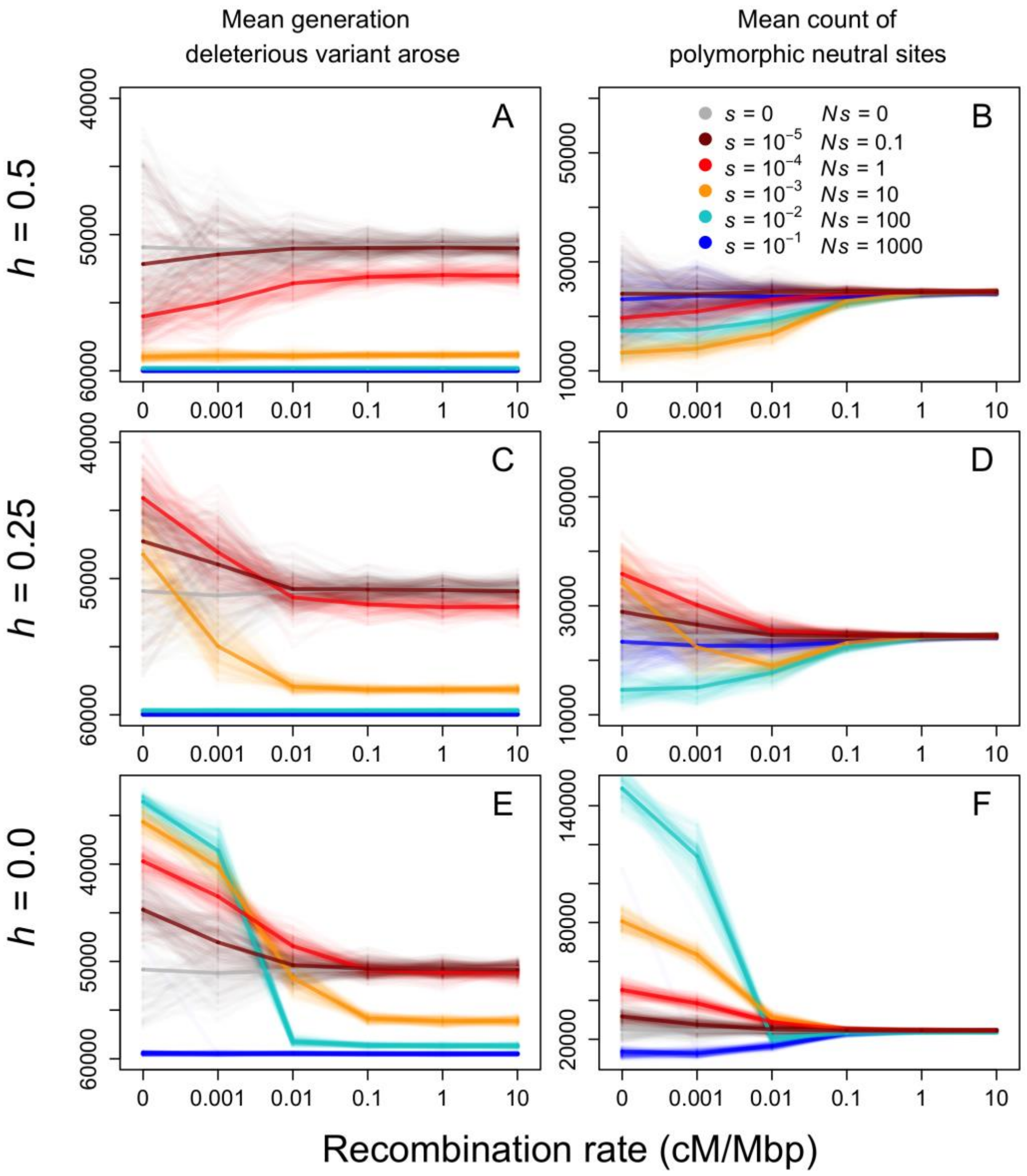

**Figure S2.** Summary of haplotypes of deleterious variants within the population for fully recessive simulations ($h = 0$). The top panel summarizes the summed length of all branches of a dendrogram from clustering individuals based on their haplotypes for deleterious heterozygote or homozygote variants, across all cases of selection coefficients and all recombination rates. Three individual replicates are chosen from the 0 recombination case, labelled as A ($s = -0.1$), B ($s = -0.001$), and C ($s = -0.00001$) in the top panel, where the dendrogram is shown on the left and individuals are in rows, while deleterious variants are in columns (number of variants present change across simulations). Clustering is performed by heatmap2 in R, where we can see a clear case of many complementary haplotypes in B, while for strongest $s$ in A, there are few and varied segregating deleterious alleles. Under weakest $s$ (C), there are large regions containing clusters of heterozygous or homozygous weakly deleterious alleles.

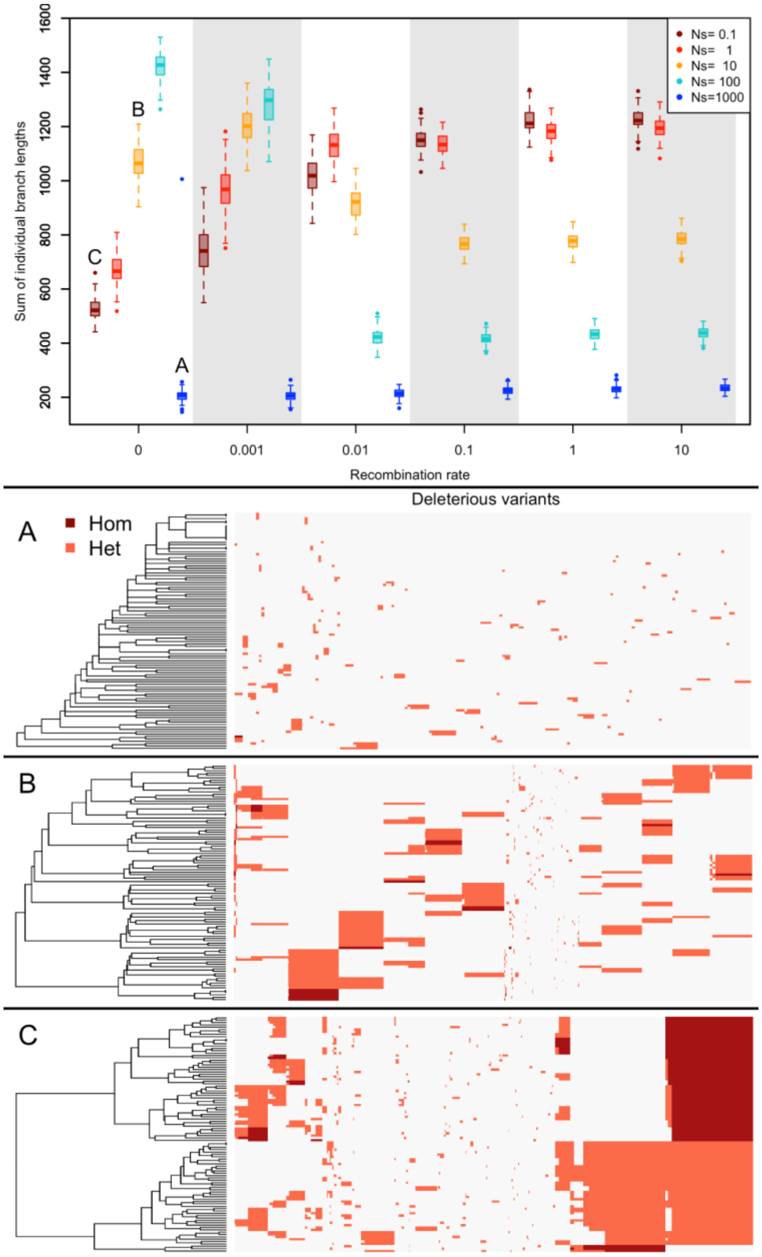

**Figure S3.** Example of 2-locus dynamics for deleterious recessive alleles (alleles denoted A and B). Panel A: With freely recombining loci, we find that the wild-type haplotype can be permanently maintained in the population and each locus is subject to purifying selection. Right panels: In the absence of recombination, the loss of the wild-type haplotype results in every individual possessing at least one deleterious allele, leading to the permanent maintenance of complementary haplotypes. Parameter values are $N = 20000$ individuals, $s = -0.01, h = 0$ and $\mu = 0.25$. Panel B: Like panel A except showing different selection intensities at the two loci ($s_A = -0.005, s_B = -0.01$).

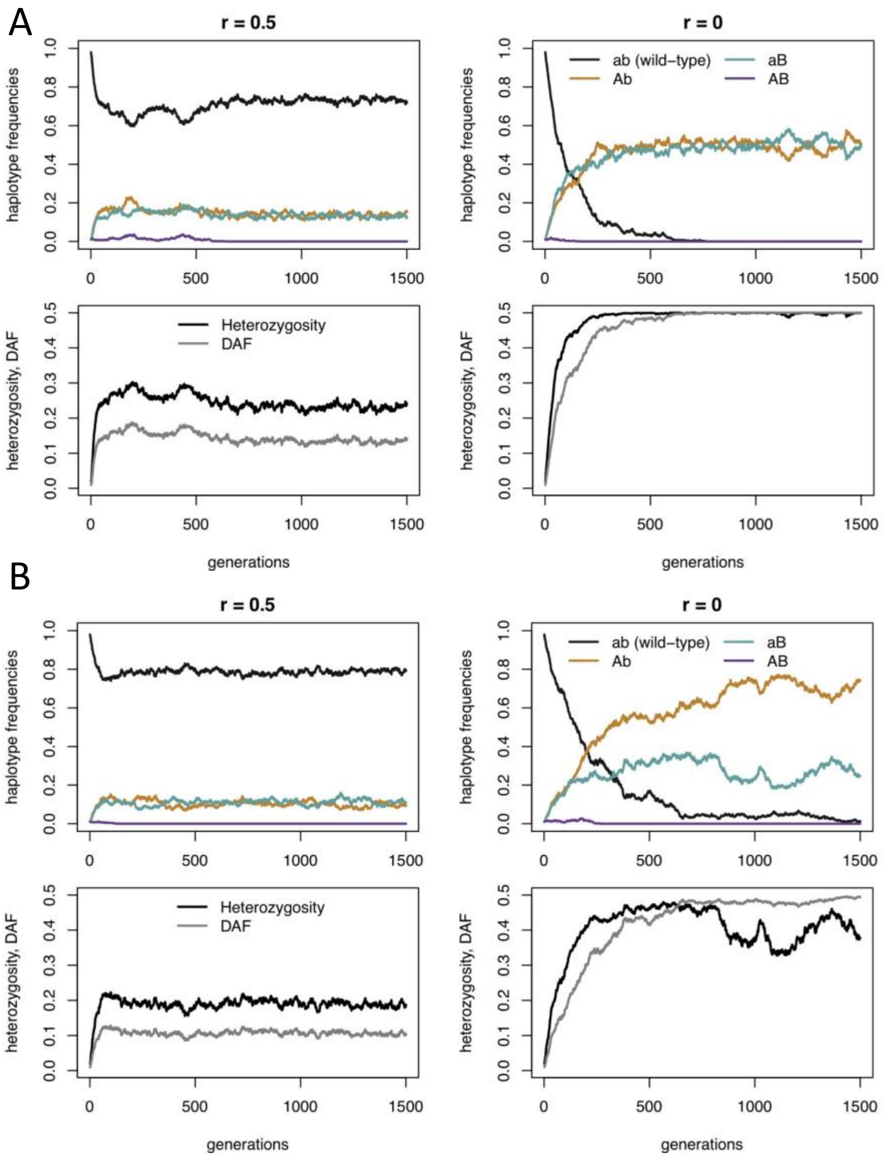

**Figure S4.** Simulations under higher deleterious mutation rate (20% of mutations are deleterious, i.e. per-bp $\mu = 2.9e^{-9}$) show that the transition from BGS to AOD occurs under different scenarios of selection coefficients (s), dominance of deleterious alleles (h), and recombination rates. Genomic diversity at neutral sites as measured by nucleotide diversity ($\pi$) and derived allele frequency ($DAF_i$) from simulated scenarios. Means across all replicates are shown in thick lines while all 100 replicates are transparent lines in the background.

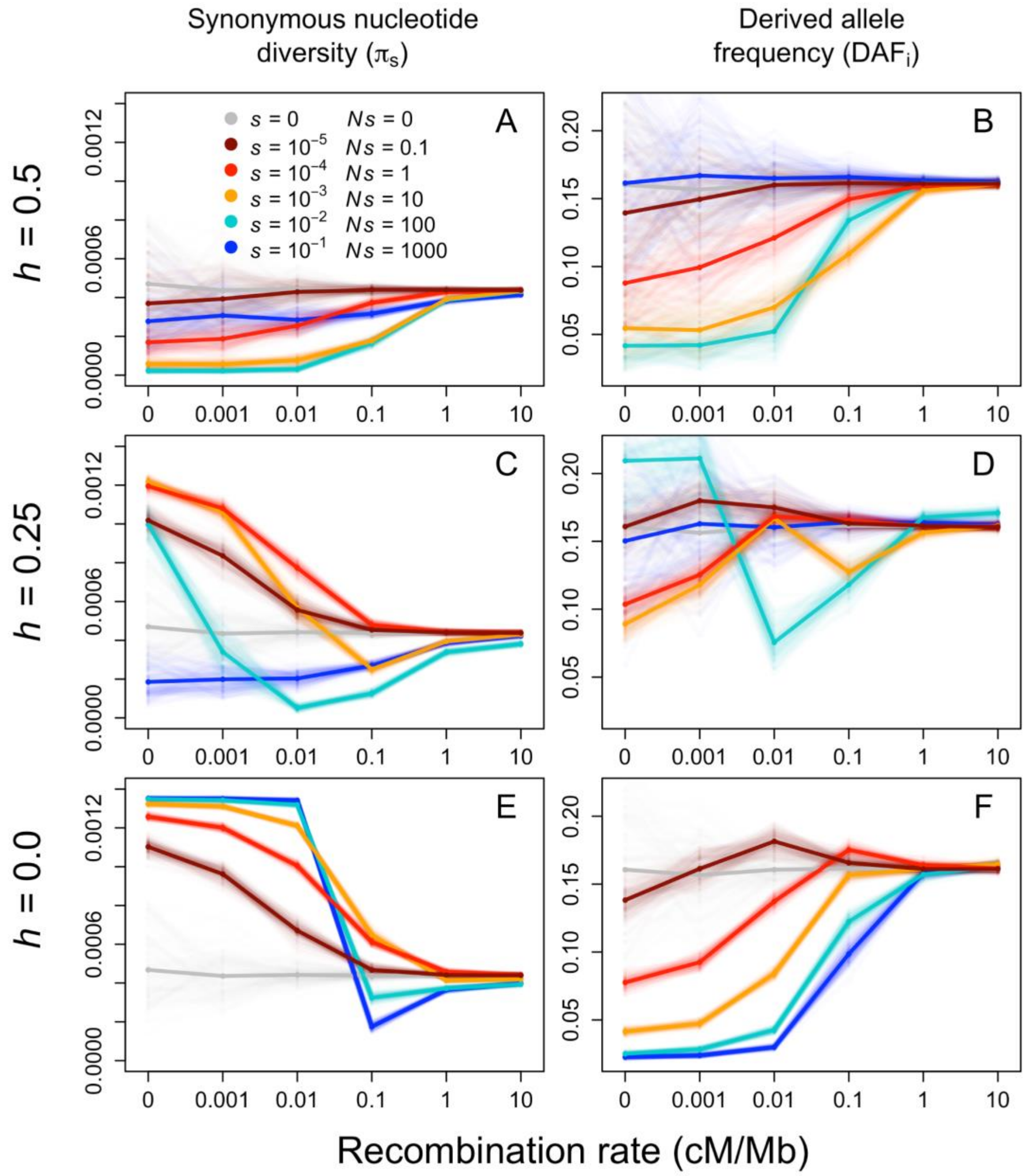

### Supplemental Item 1

#### A two-locus model without recombination

##### The Model

**We consider a two-locus, two-allele model with the alleles a and A at locus 1 and b and B at locus 2. Both derived alleles are fully recessive deleterious and their selection coefficient is denoted s. Fitness effects interact multiplicatively across loci and the fitness of all two locus genotypes is summarized by:**

| | aa | aA | AA |
|---|---|---|---|
| bb | 1 | 1 | 1 – s |
| bB | 1 | 1 | 1 – s |
| BB | 1 – s | 1 – s | $(1 - s)^2$ |

**Let $p_1$, $p_2$, $p_3$ and $p_4$ denote the frequencies of the haplotype ab, aB, Ab, and AB, respectively. We assume that there is no recombination between loci. We next derive the standard recurrence equation for the evolution of haplotype frequencies in an (infinitely) large population. Let W denote the fitness matrix where entry (i,j) gives the fitness of the two-locus genotype (i,j), where i and j specify the haplotypes:**

$$W = \begin{pmatrix} 1 & 1 & 1 & 1 \\ 1 & 1-s & 1 & 1-s \\ 1 & 1 & 1-s & 1-s \\ 1 & 1-s & 1-s & 1-2s \end{pmatrix};$$

**and let p denote the vector of haplotype frequencies**

```
p = {P1, P2, P3, P4};
```

**The marginal fitness of each haplotype is then given by**

```
Wm = W.p; MatrixForm[Wm]
```

$$\begin{pmatrix} P1 + P2 + P3 + P4 \\ P1 + P3 + P2\ (1 - s) + P4\ (1 - s) \\ P1 + P2 + P3\ (1 - s) + P4\ (1 - s) \\ P1 + P4\ (1 - 2\ s) + P2\ (1 - s) + P3\ (1 - s) \end{pmatrix}$$

**and the mean fitness of the populationd by**

```
Wbar = p.Wm;
```

**The expected haplotype frequencies p' after selection are:**

```
p' = p * Wm / Wbar;
```

**and after reproduction and mutation, denoted p'':**

$$\Upsilon = \begin{pmatrix} 0 & 0 & 0 & 0 \\ u & 0 & 0 & 0 \\ u & 0 & 0 & 0 \\ 0 & u & u & 0 \end{pmatrix};$$

```
utot = Total[ϒ]
```

```
{2 u, u, u, 0}
```

```
p'' = p' (1 - utot) + ϒ.p';
```

**where ϒ is the matrix that specifies the rate of mutation from haplotype i to j, and utot gives the total mutation rate per haplotype. For simplicity we ignore back mutations. The expected haplotype frequencies in the next generation are then:**

```
pNext = FullSimplify[p'']
```

```
{-((P1 (P1 + P2 + P3 + P4) (-1 + 2 u)) /
    ((P1 + P2 + P3 + P4)^2 - (P2^2 + P3^2 + 2 P2 P4 + 2 P4 (P3 + P4)) s)),
  (-P2 (P2 + P3 + P4 - (P2 + P4) s) (-1 + u) + P1^2 u + P1 (P2 + (P3 + P4) u)) /
   ((P1 + P2 + P3 + P4)^2 - (P2^2 + P3^2 + 2 P2 P4 + 2 P4 (P3 + P4)) s),
  (-P3 (P2 + P3 + P4 - (P3 + P4) s) (-1 + u) + P1^2 u + P1 (P3 + (P2 + P4) u)) /
   ((P1 + P2 + P3 + P4)^2 - (P2^2 + P3^2 + 2 P2 P4 + 2 P4 (P3 + P4)) s),
  (P4 (P1 + P2 + P3 + P4 - (P2 + P3 + 2 P4) s) +
      ((P2 + P3) (P1 + P2 + P3 + P4) - (P2^2 + P2 P4 + P3 (P3 + P4)) s) u) /
   ((P1 + P2 + P3 + P4)^2 - (P2^2 + P3^2 + 2 P2 P4 + 2 P4 (P3 + P4)) s)}
```

#### Equilibria

**Ignoring mutation, the equilibria are given by:**

```
Solve[p' == p, p]
```

**Solve**: Equations may not give solutions for all "solve" variables.

```
{{P1 → 1 + P4, P2 → -P4, P3 → -P4}, {P1 → 0, P2 → 1, P3 → 1, P4 → -1},
 {P1 → 1, P2 → 0, P3 → 0, P4 → 0}, {P1 → 0, P2 → 1, P3 → 0, P4 → 0},
 {P1 → 1, P2 → 0, P3 → 0, P4 → 0}, {P1 → 0, P2 → 1/2, P3 → 1/2, P4 → 0},
 {P1 → 0, P2 → 0, P3 → 1, P4 → 0}, {P1 → 0, P2 → 0, P3 → 0, P4 → 1}}
```

**We can visualize all equilibria with P4 = 0 in a 2D plot:**

```
eql = {P1, P2, P3} /. {{P1 → 1, P2 → 0, P3 → 0, P4 → 0},
    {P1 → 0, P2 → 1, P3 → 0, P4 → 0}, {P1 → 1, P2 → 0, P3 → 0, P4 → 0},
    {P1 → 0, P2 → 1/2, P3 → 1/2, P4 → 0}, {P1 → 0, P2 → 0, P3 → 1, P4 → 0}};
coords[{r_, g_, b_}] := {r / 2 + g, r Tan[Pi / 3] / 2};
cg =
  Graphics[{PointSize[Large], Disk[coords[# / Total[#]], 0.01]} & /@ eql, ImageSize → 300];
cl = Graphics[{Thick, Line[{{0, 0}, {1, 0}, {0.5, Sqrt[3] / 2}, {0, 0}}]},
   ImageSize → 300];
tg = Graphics[{Text[Style["P2 = 1", Black, Medium, Bold], {0 - 0.05, 0 - 0.05}],
    Text[Style["P3 = 1", Black, Medium, Bold], {1 + 0.05, 0 - 0.05}],
    Text[Style["P1 = 1", Black, Medium, Bold], {0.5, √3 / 2 + 0.05}]}, ImageSize → 300];
Show[cl, cg, tg, ImageSize → 300]
```

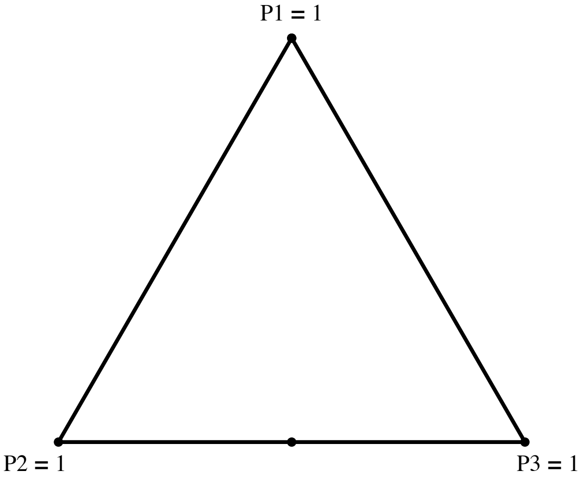


**This shows that if both loci are polymorphic and every individual carries at least one derived allele, the two haplotypes aB and Ab will be maintained at frequency 1/2. Next, we add mutations:**

```
sols = Solve[pNext == p, p]; sols /. {u → 0.001, s → 0.1}
```

```
{{P1 → 0, P2 → 0, P3 → 0, P4 → 1}, {P1 → 0, P2 → 0, P3 → 0.905132, P4 → 0.0948683},
 {P1 → 0, P2 → 0, P3 → 1.09487, P4 → -0.0948683},
 {P1 → 0, P2 → 0.905132, P3 → -6.02401 × 10^-17, P4 → 0.0948683},
 {P1 → 0, P2 → 1.09487, P3 → 6.02401 × 10^-17, P4 → -0.0948683},
 {P1 → 0, P2 → 0.490683, P3 → 0.490683, P4 → 0.0186341},
 {P1 → 0, P2 → 1.00982, P3 → 1.00982, P4 → -1.01963},
 {P1 → 1.21011, P2 → -0.11011, P3 → -0.11011, P4 → 0.0101101},
 {P1 → 0.80991, P2 → 0.0900901, P3 → 0.0900901, P4 → 0.00990991},
 {P1 → 0.98999, P2 → -0.0990216, P3 → 0.100077, P4 → 0.00895434},
 {P1 → 0.98999, P2 → 0.100077, P3 → -0.0990216, P4 → 0.00895434},
 {P1 → 0.98999, P2 → -0.0905164, P3 → 0.109481, P4 → -0.00895434},
 {P1 → 0.98999, P2 → 0.109481, P3 → -0.0905164, P4 → -0.00895434}}
```

**We now see that there is an additional feasible (internal) equilibrium now. It is given by**

```
sols[[9]] // FullSimplify
```

$$\left\{P1 \to \left(s^2(-1+u)^2 - s(-1+u)u + (-2+u)\sqrt{s^3(-1+u)^2 u}\right) \Big/ \left(s^2(-1+u)^2\right),\right.$$

$$P2 \to \frac{s(-1+u)u + \sqrt{s^3(-1+u)^2 u}}{s^2(-1+u)^2},\ P3 \to \frac{s(-1+u)u + \sqrt{s^3(-1+u)^2 u}}{s^2(-1+u)^2},$$

$$\left.P4 \to -\frac{u\left(s(-1+u) + \sqrt{s^3(-1+u)^2 u}\right)}{s^2(-1+u)^2}\right\}$$

```
Series[sols[[9]] /. {u → μ s}, {s, 0, 1}] /. {μ → u / s} // Normal // FullSimplify //
  PowerExpand // FullSimplify
```

$$\left\{P1 \to \frac{s(-1+u) + \sqrt{s}(-2+u)\sqrt{u} - u}{s(-1+u)},\ P2 \to \frac{\sqrt{s}\sqrt{u} + u}{s(-1+u)},\ P3 \to \frac{\sqrt{s}\sqrt{u} + u}{s(-1+u)},\ P4 \to \frac{u + \sqrt{s}\,u^{3/2}}{s - s\,u}\right\}$$

**Note that this solution is approximately {1-2 $\sqrt{u/s}$,$\sqrt{u/s}$,$\sqrt{u/s}$, 0} and can thus be viewed as the 2-locus version of the classical result that the derived allele frequency is $\sqrt{u/s}$ at mutation-selection balance.**

**All feasible equilibria are:**

```
sols1 = sols[[{2, 4, 6, 9}]];
```

**If we plot all feasible equilibria, we can see that the internal equilibrium leaves the state space if the mutation rate becomes to high. the internal equilibrium where all haplotypes are present, disappears and the new globally stable equilibrium is the one where haplotypes aB and Ab are maintained at roughly 50% each. This bifurcation corresponds to the transition from purifying selection to pseudo-overdominance where complementary haplotypes are maintained by selection.**

```
Manipulate[
 Quiet[eql = {P1, P2, P3} /. sols1 /. {u → MutationRate, s → SelectionCoefficient}];
 coords[{r_, g_, b_}] := {r / 2 + g, r Tan[Pi / 3] / 2};
 Quiet[cg = Graphics[
    {PointSize[Large], Disk[coords[# / Total[#]], 0.01]} & /@ eql, ImageSize → 300]];
 cl = Graphics[{Thick, Line[{{0, 0}, {1, 0}, {0.5, Sqrt[3] / 2}, {0, 0}}]},
   ImageSize → 300];
 tg = Graphics[{Text[Style["P2 = 1", Black, Medium, Bold], {0 - 0.05, 0 - 0.05}],
    Text[Style["P3 = 1", Black, Medium, Bold], {1 + 0.05, 0 - 0.05}],
    Text[Style["P1 = 1", Black, Medium, Bold], {0.5, √3 / 2 + 0.05}]}, ImageSize → 300];
 Quiet[Show[cl, cg, tg, ImageSize → 300]], {MutationRate, 0.000001, 0.0005},
 {SelectionCoefficient, 0.0001, 0.1}]
```

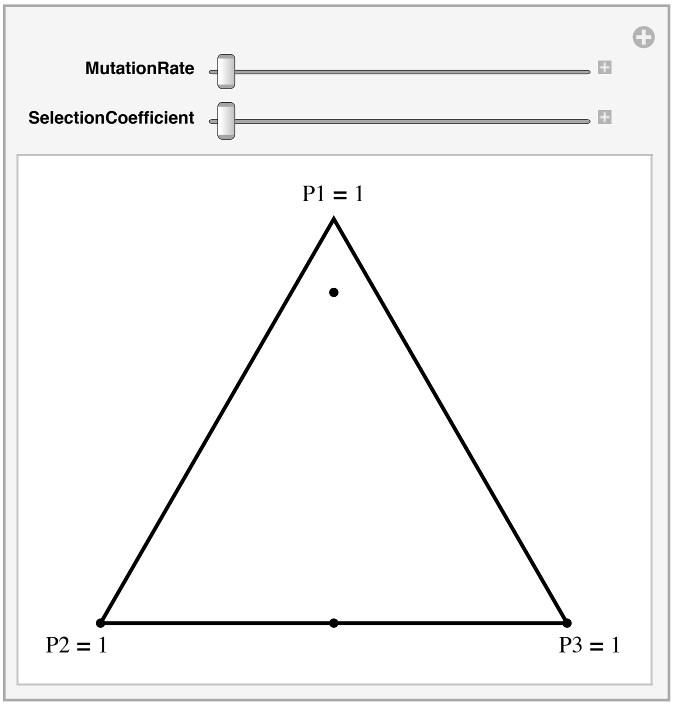


- Recombination

**We next add recombination to our calculations.**

```
LD = p[[1]] p[[4]] - p[[2]] p[[3]];
R = {r, -r, -r, r};
pNextR = p' (1 - utot) + ϒ.p' - R * LD // Simplify;
```

**To get the equilibria we need to solve the equation pNextR == p:**

```
eqn = FullSimplify[pNextR - p /. {u → s μ, r → ρ s, P4 → 1 - P1 - P2 - P3}];
```

```
Series[eqn, {s, 0, 1}] // Normal // Simplify
```

$$\{s\,(2\,P1^3 + P1\,(2 + P2^2 + P3^2 - 2\,\mu + P2\,(-2+\rho) + P3\,(-2+\rho) - \rho) + P2\,P3\,\rho + P1^2\,(-4 + 2\,P2 + 2\,P3 + \rho)),$$
$$s\,(2\,(-1+P1)\,P2^2 + P2^3 + P1\,(\mu - (-1 + P1 + P3)\,\rho) + P2\,(1 + 2\,P1^2 + P3^2 - \mu + P1\,(-3 + 2\,P3 - \rho) - P3\,(1+\rho))),\ s\,(2\,(-1+P1)\,P3^2 + P3^3 + P1\,(\mu - (-1 + P1 + P2)\,\rho) + P3\,(1 + 2\,P1^2 + P2^2 - \mu + P1\,(-3 + 2\,P2 - \rho) - P2\,(1+\rho))),$$
$$-s\,(2\,P1^3 + P2^3 + P2^2\,(-2 + P3) + P3\,(1 - 2\,P3 + P3^2 - \mu) + P1\,(3\,P2^2 + (-1+P3)\,(-2 + 3\,P3 - \rho) + P2\,(-5 + 4\,P3 - \rho)) + P1^2\,(-4 + 4\,P2 + 4\,P3 - \rho) + P2\,(1 + P3^2 - \mu - P3\,(2+\rho)))\}$$

**Unfortunately, even if we ignore second and higher-order terms in s, we do not get anything useful. We thus resort to an alternative approach.**

### A multi-locus approach

#### ■ Frequency of wild-type haplotype at mutation-selection equilibrium

**We next consider a model with n loci that are equidistantly distributed over a region of length r cM so that the approximate recombination rate between two consecutive loci is r/n. The total mutation rate is denoted U = n u, where u is the per locus mutation rate. We assume that all loci are fully recessive and deleterious with selection coefficient s. To simplify calculations, we ignore haplotypes that carry more than 1 deleterious allele, as they will be vanishingly rare at mutation-selection balance. Further, since we use a symmetric model, we can assume that the frequency of haplotypes carrying the same number of derived alleles are equal at equilibrium. We denote by $p_0$ the frequency of the wild-type haplotype with 0 mutations, and by $p_1$ the frequency of a haplotype with exactly one deleterious mutation. Straightforward calculations show that the marginal fitness of these haplotypes are**

```
w0 = 1; w1 = 1 - p1 s;
```

**and the mean fitness is**

```
wbar = 1 - n (p1) ^2 s;
```

**Furthermore, we can set**

```
p1 = (1 - p0) / n;
```

**because haplotype frequencies need to sum to 1.**

**Switching to continuous time, the evolution of the frequency of the wild-type haplotype can then be approximated by**

```
dp0 / dt == p0 (w0 - wbar) - U p0 + Sum[Sum[p1^2 (j - i) r / n, {j, i, n}], {i, 1, n}] // Simplify
```

$$\frac{dp0}{dt} == \frac{1}{6\,n^2}\left((-1+n^2)\,(-1+p0)^2\,r + 6\,n\,p0\,((-1+p0)^2\,s - n\,U)\right)$$

**where the first term corresponds to selection, the second to mutation, and the last term to recombination. For the last term we assumed that double heterozygotes produce the wild-type haplotype with probability r d/n, where the two loci are d (= j - i) sites apart and the recombination rate between two consecutive sites is r/n.**

**We next rescale all quantities with respect to selection intensity and get**

$$dp0 / dt == \frac{1}{n}\left(6\,n\,p0\,((-1+p0)^2 - n^\wedge 2\,\mu) + (-1+n^2)\,(-1+p0)^2\,\rho\right),$$

**where $\mu$ = u/s and $\rho$ = r/s. We set dp0/dt ==0 to find the equilibrium states and get**

$$sols = Solve\left[\frac{1}{n}\left(6\,n\,p0\,((-1+p0)^2 - n^\wedge 2\,\mu) + (-1+n^2)\,(-1+p0)^2\,\rho\right) == 0 == 0,\ p0\right] \text{ // Simplify;}$$

**We then keep the ratio of $\sqrt{\mu} = \sqrt{u/s}$ and $\rho$ constant and ignore second and higher-order terms in s:**

```
sols2 = Series[{p0 /. sols} /. μ → (ρ ν) ^2, {ρ, 0, 2}] // Simplify // PowerExpand // Simplify
```

$$\left\{\left\{-\frac{(-1+n^2)\,\rho}{6\,n} + O[\rho]^3,\ 1 + n\,\nu\,\rho - \frac{1}{12}\left((-1+n^2)\,\nu\right)\rho^2 + O[\rho]^3,\ 1 - n\,\nu\,\rho + \frac{1}{12}(-1+n^2)\,\nu\,\rho^2 + O[\rho]^3\right\}\right\}$$

**The relevant solution is**

```
1 - n ν ρ + 1/12 (-1 + n^2) ν ρ^2 /. {ν → μ^(1/2)/ρ}
```

$$1 - n\sqrt{\mu} + \frac{1}{12}\left(-1+n^2\right)\sqrt{\mu}\,\rho$$

**Switching back to the unscaled parameters we get**

```
1 - n Sqrt[μ] + 1/12 (-1 + n^2) Sqrt[μ] ρ /. {μ → u/s, ρ → r/s}
```

$$1 - n\sqrt{\frac{u}{s}} + \frac{\left(-1+n^2\right) r \sqrt{\frac{u}{s}}}{12\,s}$$

**It is straightforward to check that this solution is also an approximate solution of the 2-locus model with recombination above by inserting the solution and developing into a Taylor Series around s = 0.**

■ Transition to pseudo-overdominance

**We now calculate the conditions for when we expect a transition from purifying selection to pseudo-overdominance and the maintenance of complementary haplotypes. This happens when $p_0 = 0$, which is the case when**

```
Solve[1 - n Sqrt[μ] + 1/12 (-1 + n^2) Sqrt[μ] ρ == 0, ρ]
```

$$\left\{\left\{\rho \to \frac{12\left(-1+n\sqrt{\mu}\right)}{\left(-1+n^2\right)\sqrt{\mu}}\right\}\right\}$$

**Thus, if $\rho < \frac{12\left(-1+n\sqrt{\mu}\right)}{\left(-1+n^2\right)\sqrt{\mu}}$ we expect a transition to maintenance of complementary haplotypes, rather than purifying selection.**

**In the unscaled parameters, this translates to:**

```
Solve[1 - n Sqrt[u/s] + 1/12 (-1 + n^2) Sqrt[u/s] r/s == 0, r]
```

$$\left\{\left\{r \to \frac{12\,s\left(-1+n\sqrt{\frac{u}{s}}\right)}{\left(-1+n^2\right)\sqrt{\frac{u}{s}}}\right\}\right\}$$

**The parameter range for which we expect loss of the fittest haplotype is indicated in the Figure below:**

```
LogLogPlot[(12 s (-1 + n Sqrt[u/s]))/((-1 + n^2) Sqrt[u/s]) /. {n → 250, u → 2 * 10^(-8)},
  {s, 0.00000001, 0.1}, Filling → Bottom, AxesLabel → {"s", "r"}]
```

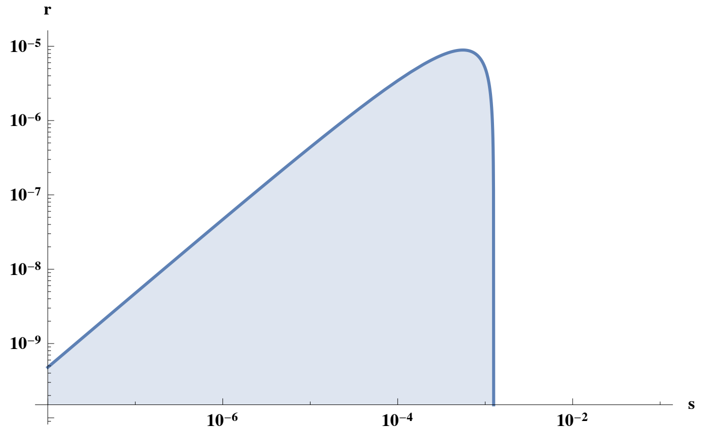


■ Equilibrium haplotype frequencies after transition to pseudo-overdominance

**We can also calculate the expected frequency of the one-mutation haplotype $p_1$. Note that even though there are n loci that can**

**potentially segregate, not all of these will be polymorphic at equilibrium. As soon as the wild-type haplotype is lost, i.e., $p_0 = 0$, the dynamics change from purging of deleterious mutations to maintenance of k distinct haploids, each at frequency 1/k. This critical k is simply the solution of**

$$\text{sols} = \text{Solve}\left[1 - k\sqrt{\mu} + \frac{1}{12}\left(-1+k^2\right)\sqrt{\mu}\,\rho == 0,\ k\right]$$

$$\left\{\left\{k \to \frac{6\left(\sqrt{\mu} - \frac{1}{6}\sqrt{36\,\mu - 12\sqrt{\mu}\,\rho + \mu\,\rho^2}\right)}{\sqrt{\mu}\,\rho}\right\}, \left\{k \to \frac{6\left(\sqrt{\mu} + \frac{1}{6}\sqrt{36\,\mu - 12\sqrt{\mu}\,\rho + \mu\,\rho^2}\right)}{\sqrt{\mu}\,\rho}\right\}\right\}$$

**The larger of the two values is the relevant solution. The expected frequency of the one-mutation haplotypes is then:**

$$\left(1/k\ /.\ k \to \frac{1}{\sqrt{\mu}\,\rho} 6\left(\sqrt{\mu} + \frac{1}{6}\sqrt{36\,\mu - 12\sqrt{\mu}\,\rho + \mu\,\rho^2}\right)\right)\ /.\ \{\mu \to u/s,\ \rho \to r/s\}$$

$$\frac{r\sqrt{\frac{u}{s}}}{6\,s\left(\sqrt{\frac{u}{s}} + \frac{1}{6}\sqrt{\frac{r^2 u}{s^3} + \frac{36\,u}{s} - \frac{12\,r\sqrt{\frac{u}{s}}}{s}}\right)}$$

**In the absence of recombination this is simply:**

$$\text{sols} = \text{Solve}\left[1 - k\sqrt{\mu} == 0,\ k\right]$$

$$\left\{\left\{k \to \frac{1}{\sqrt{\mu}}\right\}\right\}$$

**Thus, the frequency of the derived allele at each locus decreases with increasing selection coefficient.**

$$\text{LogLogPlot}\left[\sqrt{u/s}\ /.\ \{u \text{->} 2.9 * 10^{\wedge}(-8)\},\ \{s, 0, 1\},\ \text{AxesLabel} \to \{\text{"s"}, \text{"p}_1\text{"}\}\right]$$

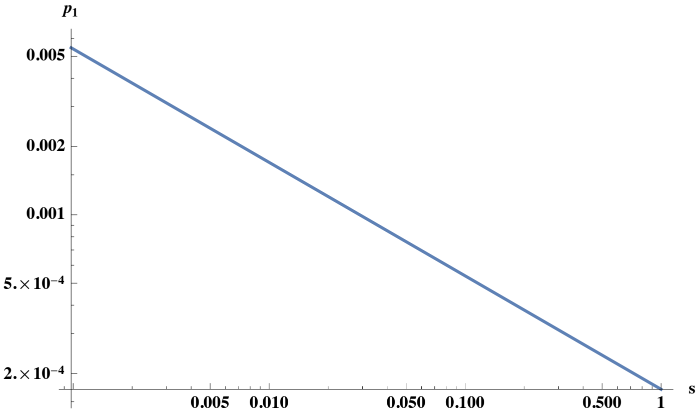

### Supplemental Item 2 of Human Analyses

Fanny Pouyet

August 2019

#### 1 Item 2A: $\pi$ scans

Same as panel A of Figure 4 but for the 22 autosomes.

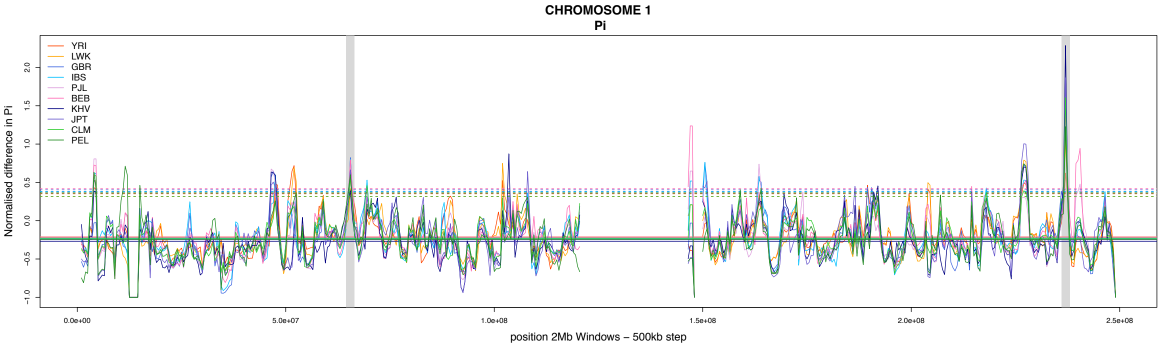
CHROMOSOME 1
Pi
YRI
LWK
GBR
IBS
PJL
BEB
KHV
JPT
CLM
PEL
Normalised difference in Pi
position 2Mb Windows – 500kb step

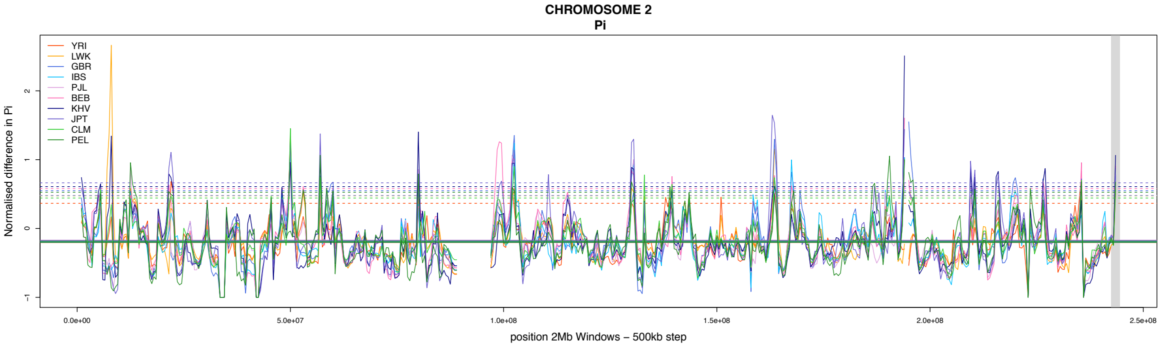
CHROMOSOME 2
Pi
YRI
LWK
GBR
IBS
PJL
BEB
KHV
JPT
CLM
PEL
Normalised difference in Pi
position 2Mb Windows – 500kb step

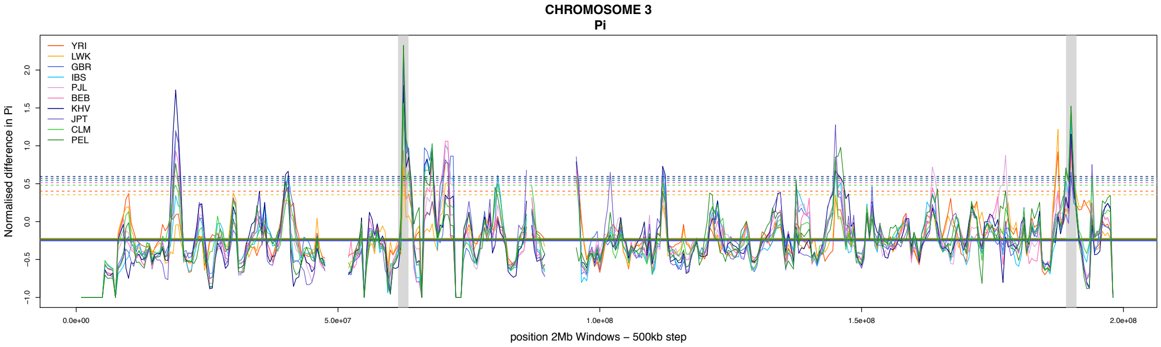
CHROMOSOME 3
Pi
YRI
LWK
GBR
IBS
PJL
BEB
KHV
JPT
CLM
PEL
Normalised difference in Pi
position 2Mb Windows – 500kb step

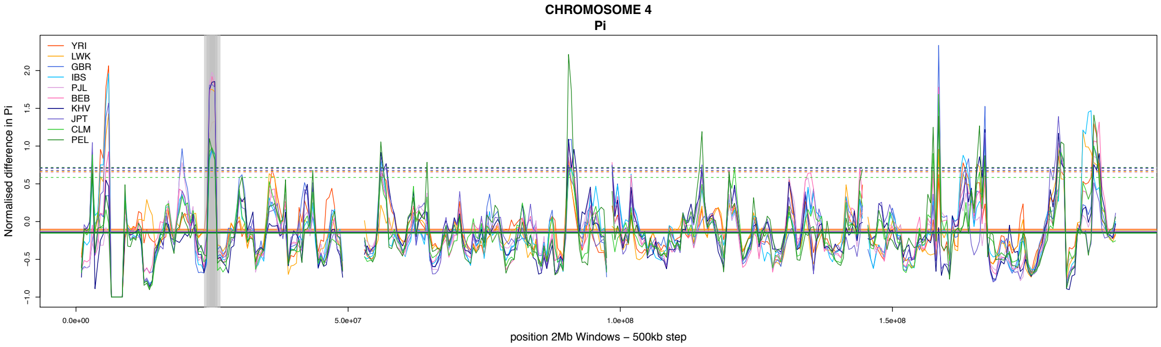
CHROMOSOME 4
Pi
YRI
LWK
GBR
IBS
PJL
BEB
KHV
JPT
CLM
PEL
Normalised difference in Pi
position 2Mb Windows – 500kb step

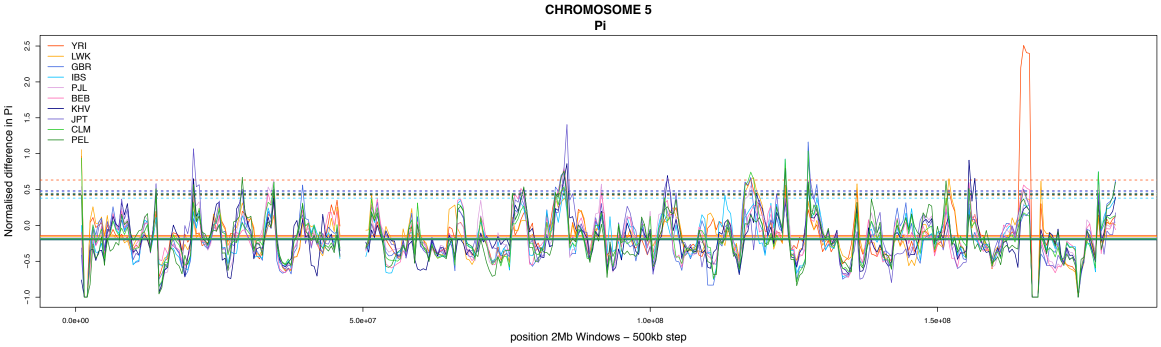
CHROMOSOME 5
Pi
YRI
LWK
GBR
IBS
PJL
BEB
KHV
JPT
CLM
PEL
Normalised difference in Pi
position 2Mb Windows – 500kb step

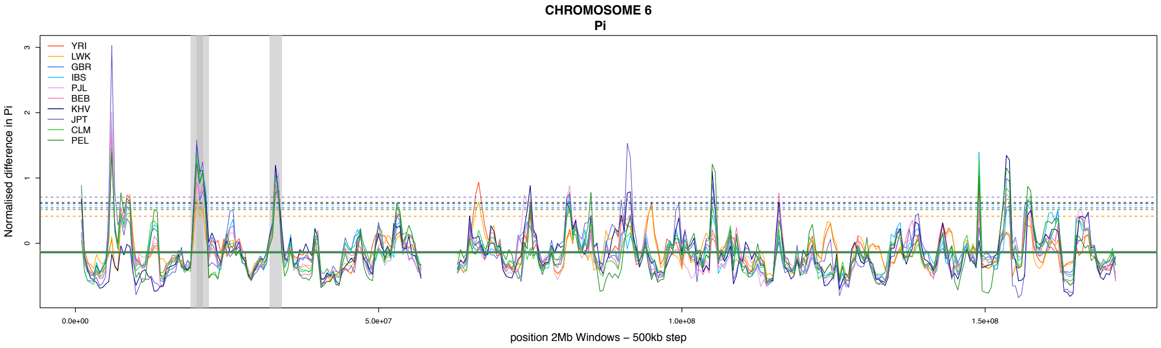
CHROMOSOME 6
Pi
YRI
LWK
GBR
IBS
PJL
BEB
KHV
JPT
CLM
PEL
Normalised difference in Pi
position 2Mb Windows – 500kb step

**CHROMOSOME 7**
**Pi**

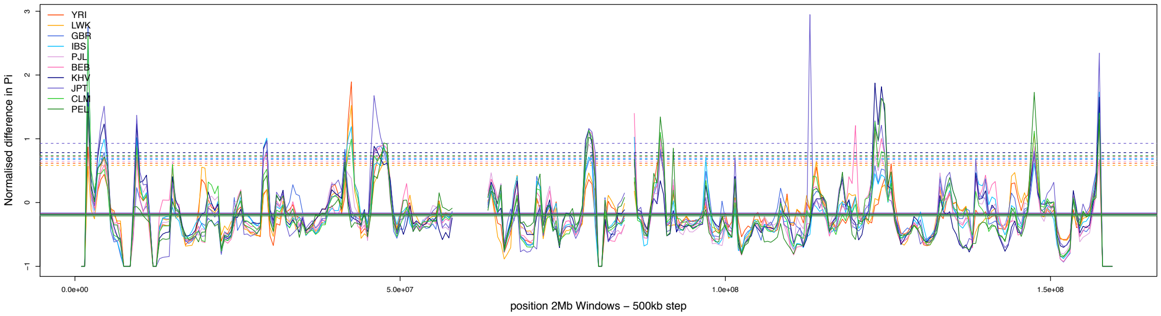


**CHROMOSOME 8**
**Pi**

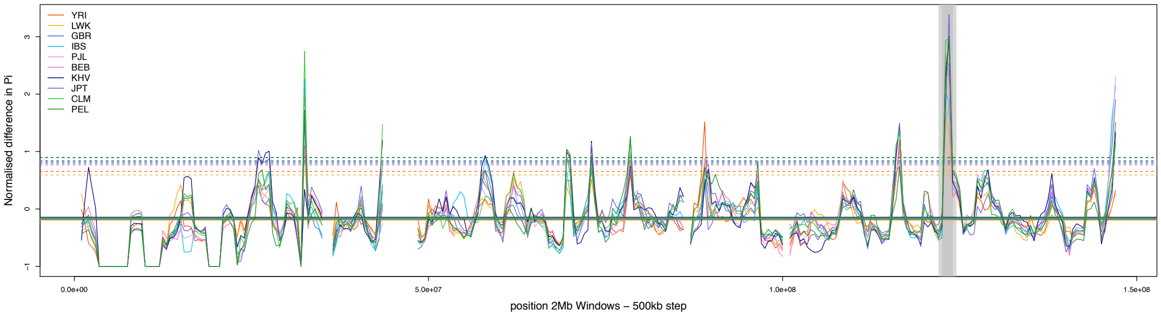


**CHROMOSOME 9**
**Pi**

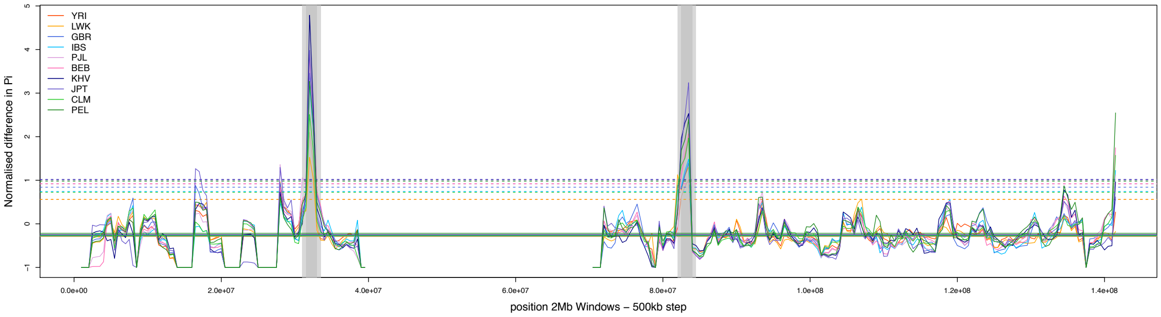


**CHROMOSOME 10**
**Pi**

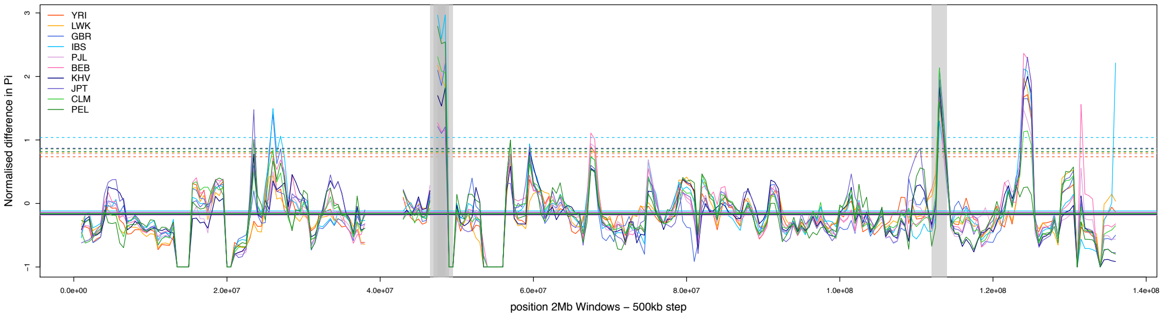


**CHROMOSOME 11**
**Pi**

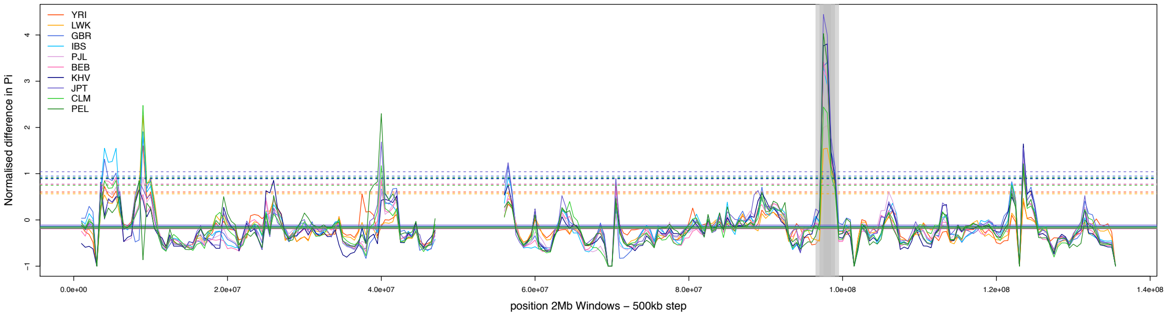


**CHROMOSOME 12**
**Pi**

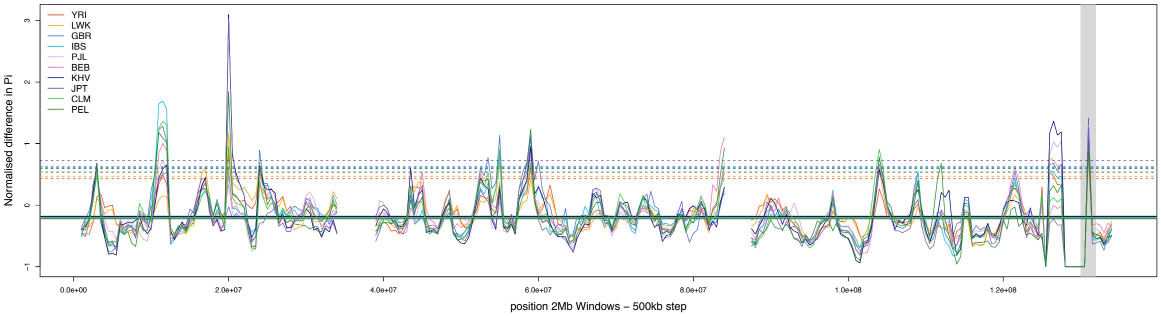

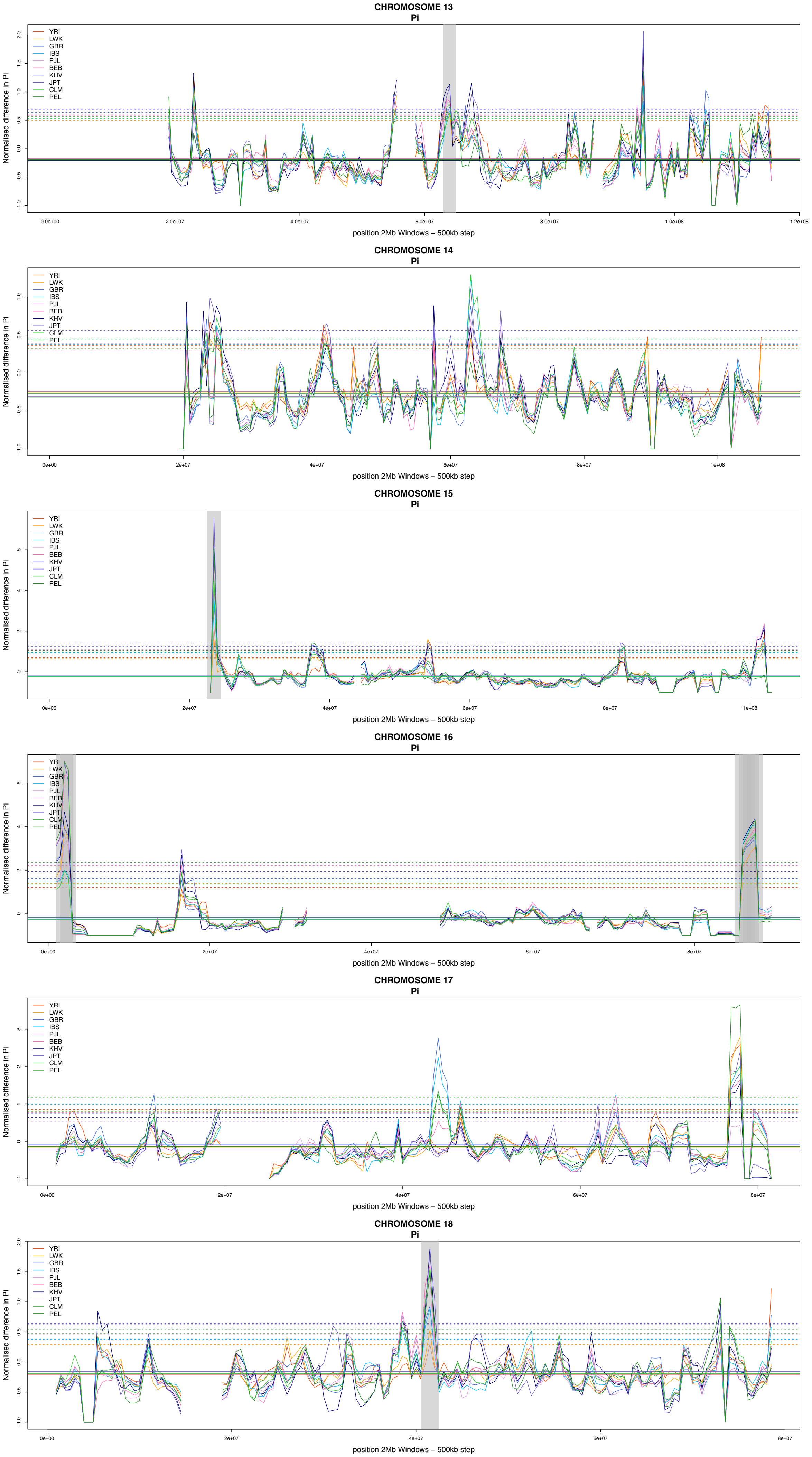

CHROMOSOME 13
Pi
YRI
LWK
GBR
IBS
PJL
BEB
KHV
JPT
CLM
PEL
Normalised difference in Pi
position 2Mb Windows – 500kb step
CHROMOSOME 14
Pi
YRI
LWK
GBR
IBS
PJL
BEB
KHV
JPT
CLM
PEL
Normalised difference in Pi
position 2Mb Windows – 500kb step
CHROMOSOME 15
Pi
YRI
LWK
GBR
IBS
PJL
BEB
KHV
JPT
CLM
PEL
Normalised difference in Pi
position 2Mb Windows – 500kb step
CHROMOSOME 16
Pi
YRI
LWK
GBR
IBS
PJL
BEB
KHV
JPT
CLM
PEL
Normalised difference in Pi
position 2Mb Windows – 500kb step
CHROMOSOME 17
Pi
YRI
LWK
GBR
IBS
PJL
BEB
KHV
JPT
CLM
PEL
Normalised difference in Pi
position 2Mb Windows – 500kb step
CHROMOSOME 18
Pi
YRI
LWK
GBR
IBS
PJL
BEB
KHV
JPT
CLM
PEL
Normalised difference in Pi
position 2Mb Windows – 500kb step

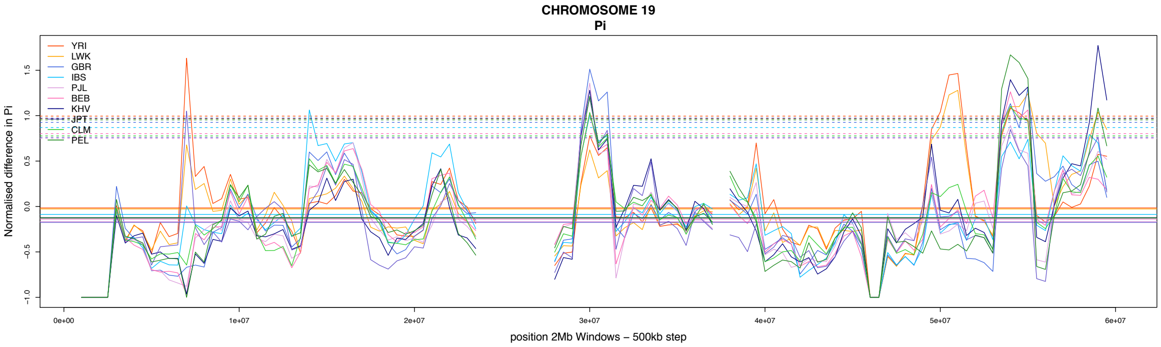
CHROMOSOME 19
Pi
YRI
LWK
GBR
IBS
PJL
BEB
KHV
JPT
CLM
PEL
Normalised difference in Pi
-1.0
-0.5
0.0
0.5
1.0
1.5
0e+00
1e+07
2e+07
3e+07
4e+07
5e+07
6e+07
position 2Mb Windows – 500kb step

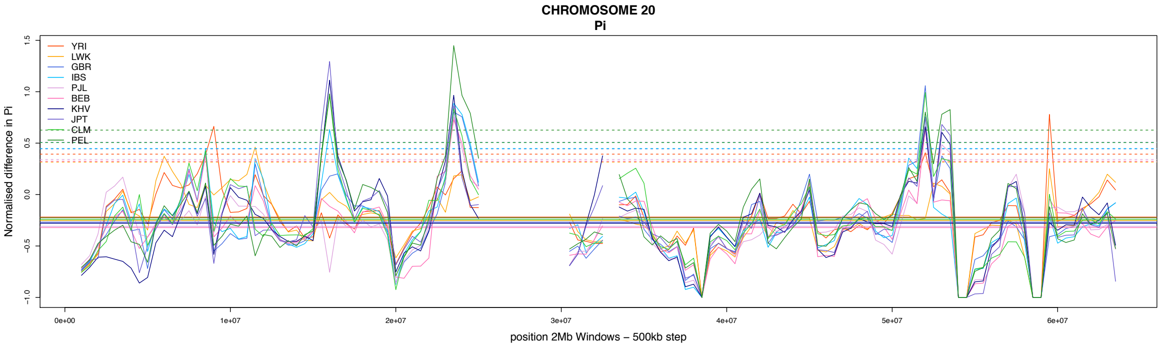
CHROMOSOME 20
Pi
YRI
LWK
GBR
IBS
PJL
BEB
KHV
JPT
CLM
PEL
Normalised difference in Pi
-1.0
-0.5
0.0
0.5
1.0
1.5
0e+00
1e+07
2e+07
3e+07
4e+07
5e+07
6e+07
position 2Mb Windows – 500kb step

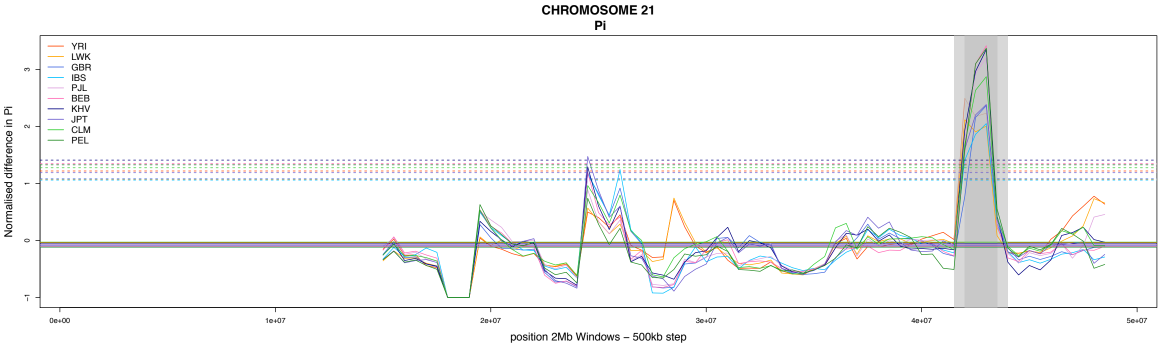
CHROMOSOME 21
Pi
YRI
LWK
GBR
IBS
PJL
BEB
KHV
JPT
CLM
PEL
Normalised difference in Pi
-1
0
1
2
3
0e+00
1e+07
2e+07
3e+07
4e+07
5e+07
position 2Mb Windows – 500kb step

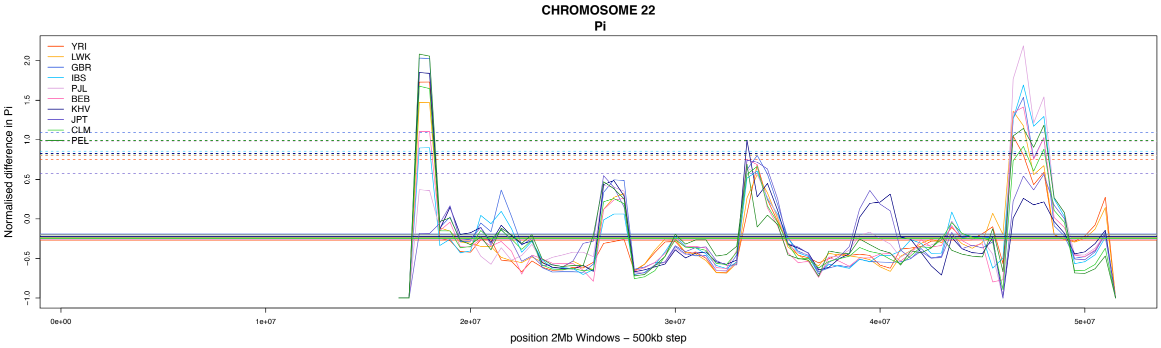
CHROMOSOME 22
Pi
YRI
LWK
GBR
IBS
PJL
BEB
KHV
JPT
CLM
PEL
Normalised difference in Pi
-1.0
-0.5
0.0
0.5
1.0
1.5
2.0
0e+00
1e+07
2e+07
3e+07
4e+07
5e+07
position 2Mb Windows – 500kb step

#### 2 Item 2B : SFS of the AOD candidate windows

Same as panel B of Figure 4 but for the 21 candidate regions for AOD. There are 4 candidate regions per page, each one consisting of 5 column and 2 rows. Populations are ordered as following: the first column consist of 2 African populations (YRI and LWK), the second column, Europeans (GBR and IBS), third column are Southern-Asian (BEB and PJL), fourth are East Asian (KHV and JPT) then Amerindians (CLM and PEL).

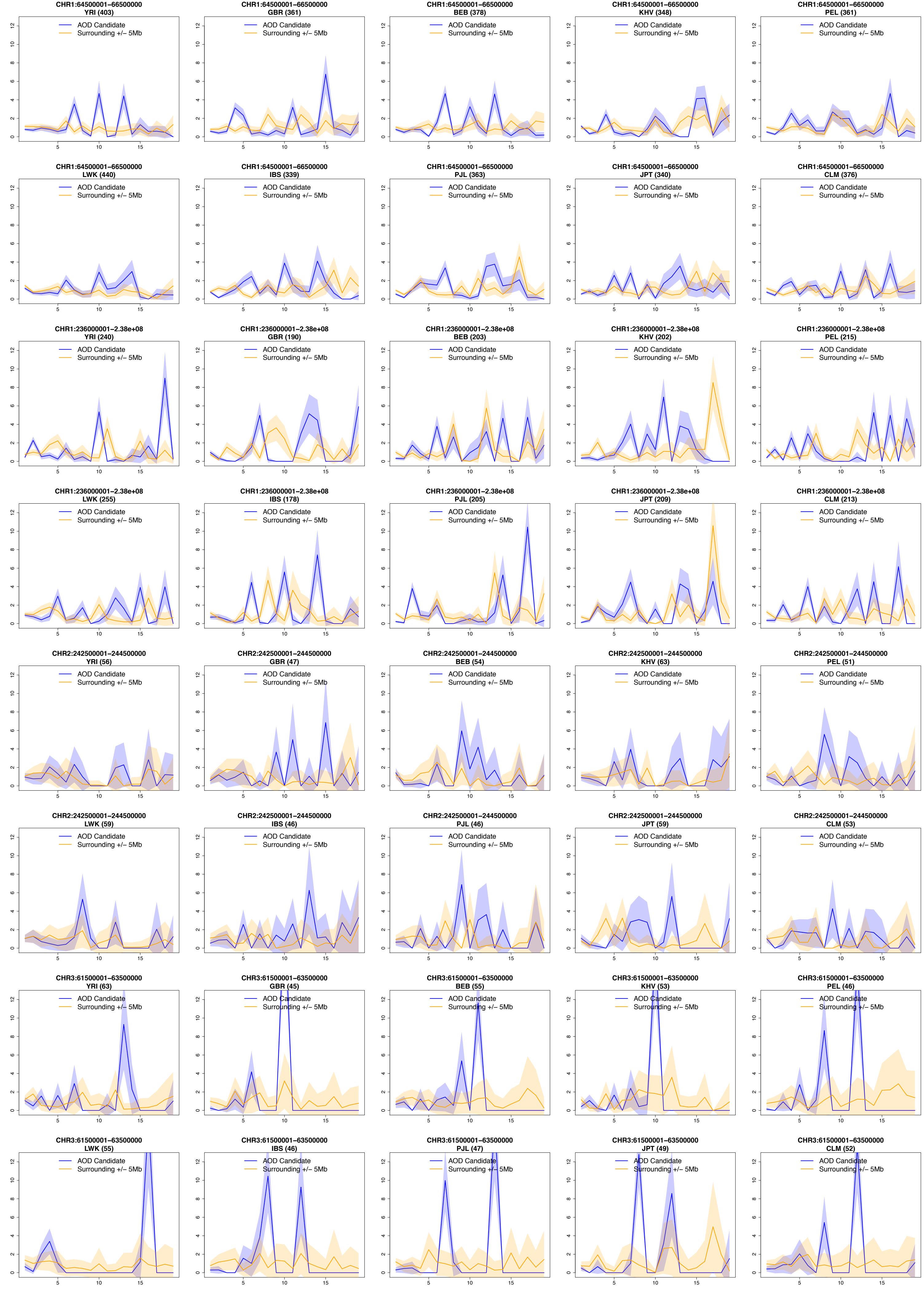
CHR1:64500001–66500000
YRI (403)
AOD Candidate
Surrounding +/– 5Mb
CHR1:64500001–66500000
GBR (361)
AOD Candidate
Surrounding +/– 5Mb
CHR1:64500001–66500000
BEB (378)
AOD Candidate
Surrounding +/– 5Mb
CHR1:64500001–66500000
KHV (348)
AOD Candidate
Surrounding +/– 5Mb
CHR1:64500001–66500000
PEL (361)
AOD Candidate
Surrounding +/– 5Mb
CHR1:64500001–66500000
LWK (440)
AOD Candidate
Surrounding +/– 5Mb
CHR1:64500001–66500000
IBS (339)
AOD Candidate
Surrounding +/– 5Mb
CHR1:64500001–66500000
PJL (363)
AOD Candidate
Surrounding +/– 5Mb
CHR1:64500001–66500000
JPT (340)
AOD Candidate
Surrounding +/– 5Mb
CHR1:64500001–66500000
CLM (376)
AOD Candidate
Surrounding +/– 5Mb
CHR1:236000001–2.38e+08
YRI (240)
AOD Candidate
Surrounding +/– 5Mb
CHR1:236000001–2.38e+08
GBR (190)
AOD Candidate
Surrounding +/– 5Mb
CHR1:236000001–2.38e+08
BEB (203)
AOD Candidate
Surrounding +/– 5Mb
CHR1:236000001–2.38e+08
KHV (202)
AOD Candidate
Surrounding +/– 5Mb
CHR1:236000001–2.38e+08
PEL (215)
AOD Candidate
Surrounding +/– 5Mb
CHR1:236000001–2.38e+08
LWK (255)
AOD Candidate
Surrounding +/– 5Mb
CHR1:236000001–2.38e+08
IBS (178)
AOD Candidate
Surrounding +/– 5Mb
CHR1:236000001–2.38e+08
PJL (205)
AOD Candidate
Surrounding +/– 5Mb
CHR1:236000001–2.38e+08
JPT (209)
AOD Candidate
Surrounding +/– 5Mb
CHR1:236000001–2.38e+08
CLM (213)
AOD Candidate
Surrounding +/– 5Mb
CHR2:242500001–244500000
YRI (56)
AOD Candidate
Surrounding +/– 5Mb
CHR2:242500001–244500000
GBR (47)
AOD Candidate
Surrounding +/– 5Mb
CHR2:242500001–244500000
BEB (54)
AOD Candidate
Surrounding +/– 5Mb
CHR2:242500001–244500000
KHV (63)
AOD Candidate
Surrounding +/– 5Mb
CHR2:242500001–244500000
PEL (51)
AOD Candidate
Surrounding +/– 5Mb
CHR2:242500001–244500000
LWK (59)
AOD Candidate
Surrounding +/– 5Mb
CHR2:242500001–244500000
IBS (46)
AOD Candidate
Surrounding +/– 5Mb
CHR2:242500001–244500000
PJL (46)
AOD Candidate
Surrounding +/– 5Mb
CHR2:242500001–244500000
JPT (59)
AOD Candidate
Surrounding +/– 5Mb
CHR2:242500001–244500000
CLM (53)
AOD Candidate
Surrounding +/– 5Mb
CHR3:61500001–63500000
YRI (63)
AOD Candidate
Surrounding +/– 5Mb
CHR3:61500001–63500000
GBR (45)
AOD Candidate
Surrounding +/– 5Mb
CHR3:61500001–63500000
BEB (55)
AOD Candidate
Surrounding +/– 5Mb
CHR3:61500001–63500000
KHV (53)
AOD Candidate
Surrounding +/– 5Mb
CHR3:61500001–63500000
PEL (46)
AOD Candidate
Surrounding +/– 5Mb
CHR3:61500001–63500000
LWK (55)
AOD Candidate
Surrounding +/– 5Mb
CHR3:61500001–63500000
IBS (46)
AOD Candidate
Surrounding +/– 5Mb
CHR3:61500001–63500000
PJL (47)
AOD Candidate
Surrounding +/– 5Mb
CHR3:61500001–63500000
JPT (49)
AOD Candidate
Surrounding +/– 5Mb
CHR3:61500001–63500000
CLM (52)
AOD Candidate
Surrounding +/– 5Mb

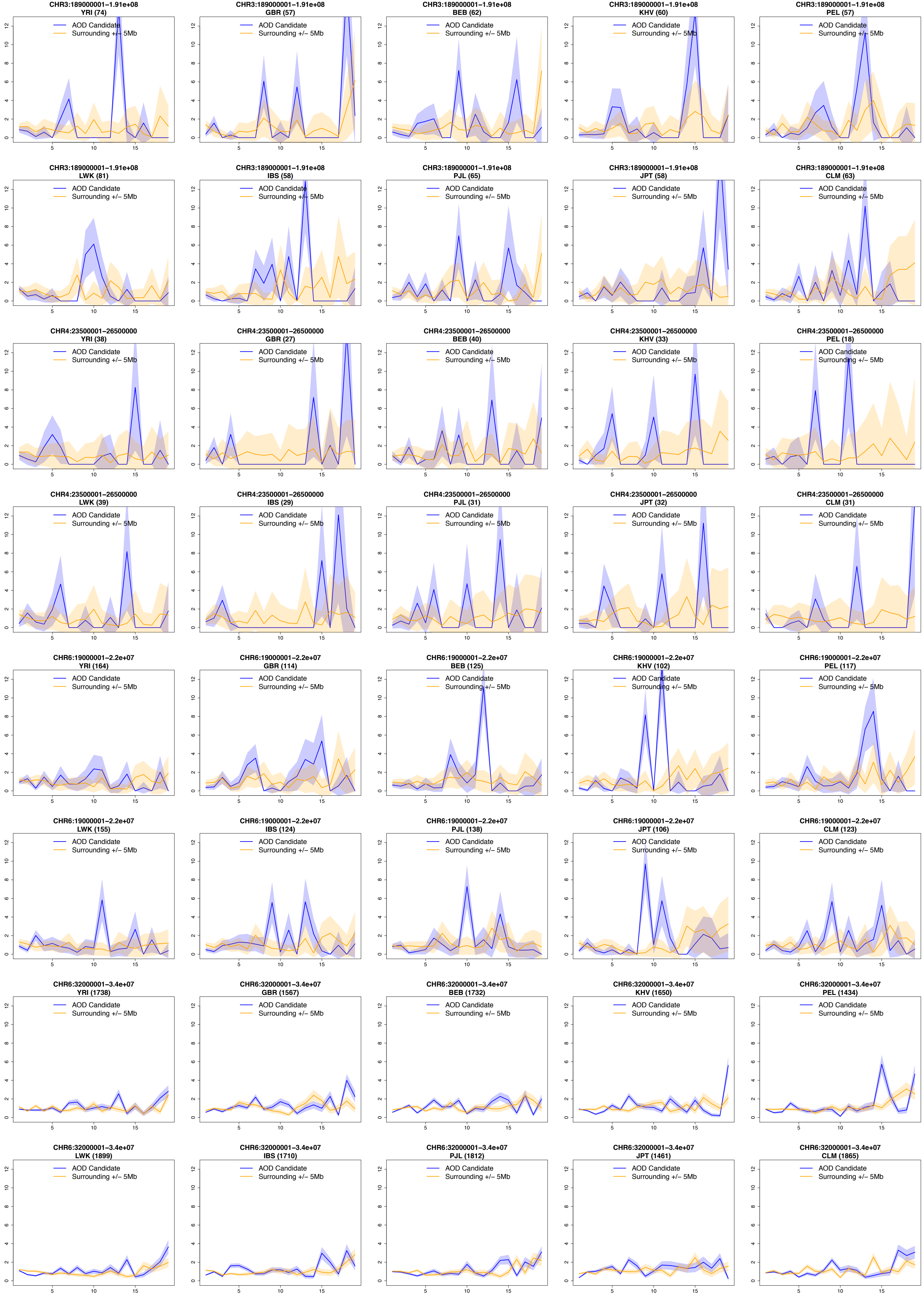
CHR3:189000001–1.91e+08
YRI (74)
AOD Candidate
Surrounding +/– 5Mb
CHR3:189000001–1.91e+08
GBR (57)
AOD Candidate
Surrounding +/– 5Mb
CHR3:189000001–1.91e+08
BEB (62)
AOD Candidate
Surrounding +/– 5Mb
CHR3:189000001–1.91e+08
KHV (60)
AOD Candidate
Surrounding +/– 5Mb
CHR3:189000001–1.91e+08
PEL (57)
AOD Candidate
Surrounding +/– 5Mb
CHR3:189000001–1.91e+08
LWK (81)
AOD Candidate
Surrounding +/– 5Mb
CHR3:189000001–1.91e+08
IBS (58)
AOD Candidate
Surrounding +/– 5Mb
CHR3:189000001–1.91e+08
PJL (65)
AOD Candidate
Surrounding +/– 5Mb
CHR3:189000001–1.91e+08
JPT (58)
AOD Candidate
Surrounding +/– 5Mb
CHR3:189000001–1.91e+08
CLM (63)
AOD Candidate
Surrounding +/– 5Mb
CHR4:23500001–26500000
YRI (38)
AOD Candidate
Surrounding +/– 5Mb
CHR4:23500001–26500000
GBR (27)
AOD Candidate
Surrounding +/– 5Mb
CHR4:23500001–26500000
BEB (40)
AOD Candidate
Surrounding +/– 5Mb
CHR4:23500001–26500000
KHV (33)
AOD Candidate
Surrounding +/– 5Mb
CHR4:23500001–26500000
PEL (18)
AOD Candidate
Surrounding +/– 5Mb
CHR4:23500001–26500000
LWK (39)
AOD Candidate
Surrounding +/– 5Mb
CHR4:23500001–26500000
IBS (29)
AOD Candidate
Surrounding +/– 5Mb
CHR4:23500001–26500000
PJL (31)
AOD Candidate
Surrounding +/– 5Mb
CHR4:23500001–26500000
JPT (32)
AOD Candidate
Surrounding +/– 5Mb
CHR4:23500001–26500000
CLM (31)
AOD Candidate
Surrounding +/– 5Mb
CHR6:19000001–2.2e+07
YRI (164)
AOD Candidate
Surrounding +/– 5Mb
CHR6:19000001–2.2e+07
GBR (114)
AOD Candidate
Surrounding +/– 5Mb
CHR6:19000001–2.2e+07
BEB (125)
AOD Candidate
Surrounding +/– 5Mb
CHR6:19000001–2.2e+07
KHV (102)
AOD Candidate
Surrounding +/– 5Mb
CHR6:19000001–2.2e+07
PEL (117)
AOD Candidate
Surrounding +/– 5Mb
CHR6:19000001–2.2e+07
LWK (155)
AOD Candidate
Surrounding +/– 5Mb
CHR6:19000001–2.2e+07
IBS (124)
AOD Candidate
Surrounding +/– 5Mb
CHR6:19000001–2.2e+07
PJL (138)
AOD Candidate
Surrounding +/– 5Mb
CHR6:19000001–2.2e+07
JPT (106)
AOD Candidate
Surrounding +/– 5Mb
CHR6:19000001–2.2e+07
CLM (123)
AOD Candidate
Surrounding +/– 5Mb
CHR6:32000001–3.4e+07
YRI (1738)
AOD Candidate
Surrounding +/– 5Mb
CHR6:32000001–3.4e+07
GBR (1567)
AOD Candidate
Surrounding +/– 5Mb
CHR6:32000001–3.4e+07
BEB (1732)
AOD Candidate
Surrounding +/– 5Mb
CHR6:32000001–3.4e+07
KHV (1650)
AOD Candidate
Surrounding +/– 5Mb
CHR6:32000001–3.4e+07
PEL (1434)
AOD Candidate
Surrounding +/– 5Mb
CHR6:32000001–3.4e+07
LWK (1899)
AOD Candidate
Surrounding +/– 5Mb
CHR6:32000001–3.4e+07
IBS (1710)
AOD Candidate
Surrounding +/– 5Mb
CHR6:32000001–3.4e+07
PJL (1812)
AOD Candidate
Surrounding +/– 5Mb
CHR6:32000001–3.4e+07
JPT (1461)
AOD Candidate
Surrounding +/– 5Mb
CHR6:32000001–3.4e+07
CLM (1865)
AOD Candidate
Surrounding +/– 5Mb

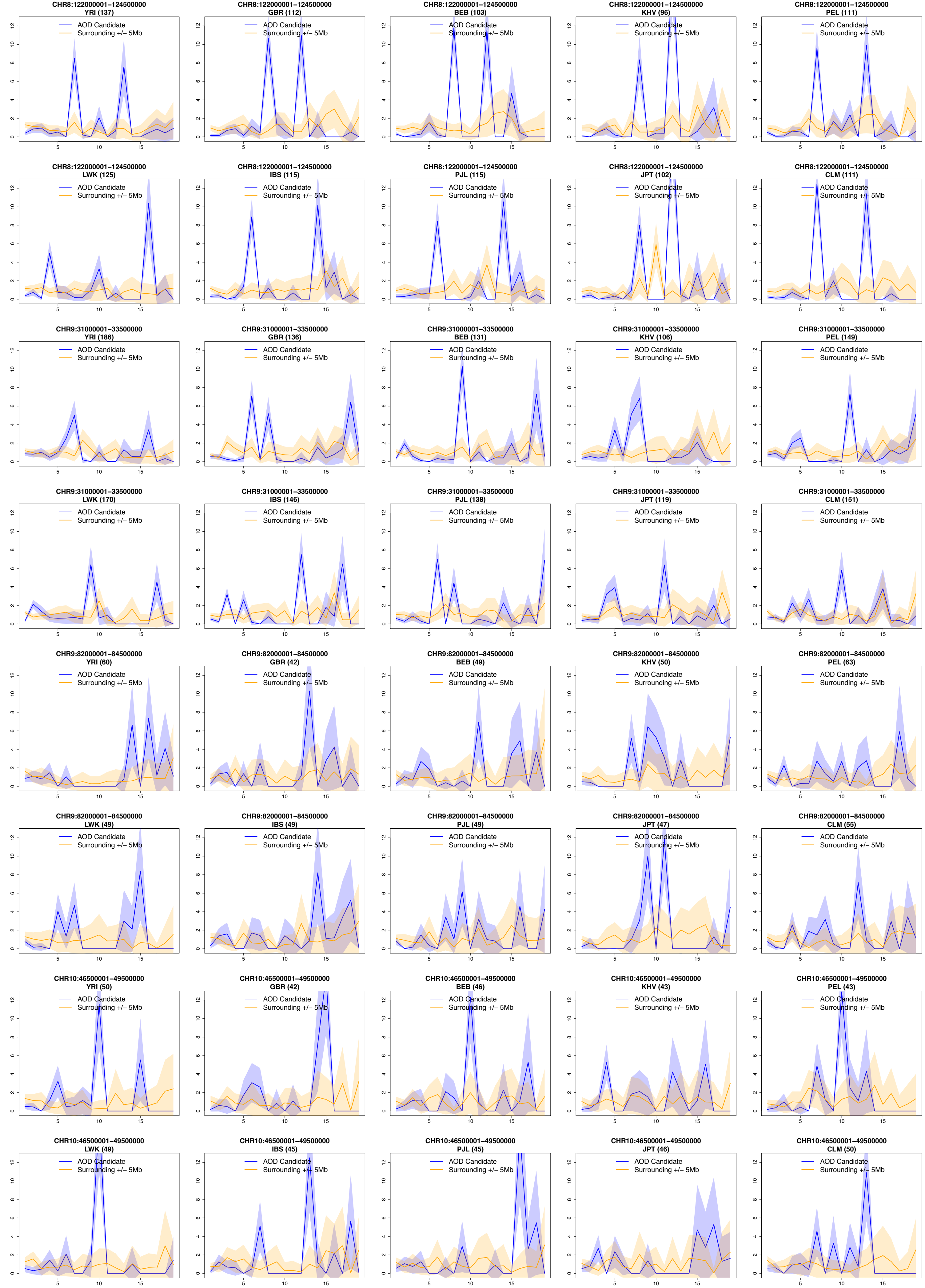
CHR8:122000001–124500000
YRI (137)
CHR8:122000001–124500000
GBR (112)
CHR8:122000001–124500000
BEB (103)
CHR8:122000001–124500000
KHV (96)
CHR8:122000001–124500000
PEL (111)
CHR8:122000001–124500000
LWK (125)
CHR8:122000001–124500000
IBS (115)
CHR8:122000001–124500000
PJL (115)
CHR8:122000001–124500000
JPT (102)
CHR8:122000001–124500000
CLM (111)
CHR9:31000001–33500000
YRI (186)
CHR9:31000001–33500000
GBR (136)
CHR9:31000001–33500000
BEB (131)
CHR9:31000001–33500000
KHV (106)
CHR9:31000001–33500000
PEL (149)
CHR9:31000001–33500000
LWK (170)
CHR9:31000001–33500000
IBS (146)
CHR9:31000001–33500000
PJL (138)
CHR9:31000001–33500000
JPT (119)
CHR9:31000001–33500000
CLM (151)
CHR9:82000001–84500000
YRI (60)
CHR9:82000001–84500000
GBR (42)
CHR9:82000001–84500000
BEB (49)
CHR9:82000001–84500000
KHV (50)
CHR9:82000001–84500000
PEL (63)
CHR9:82000001–84500000
LWK (49)
CHR9:82000001–84500000
IBS (49)
CHR9:82000001–84500000
PJL (49)
CHR9:82000001–84500000
JPT (47)
CHR9:82000001–84500000
CLM (55)
CHR10:46500001–49500000
YRI (50)
CHR10:46500001–49500000
GBR (42)
CHR10:46500001–49500000
BEB (46)
CHR10:46500001–49500000
KHV (43)
CHR10:46500001–49500000
PEL (43)
CHR10:46500001–49500000
LWK (49)
CHR10:46500001–49500000
IBS (45)
CHR10:46500001–49500000
PJL (45)
CHR10:46500001–49500000
JPT (46)
CHR10:46500001–49500000
CLM (50)
AOD Candidate
Surrounding +/− 5Mb

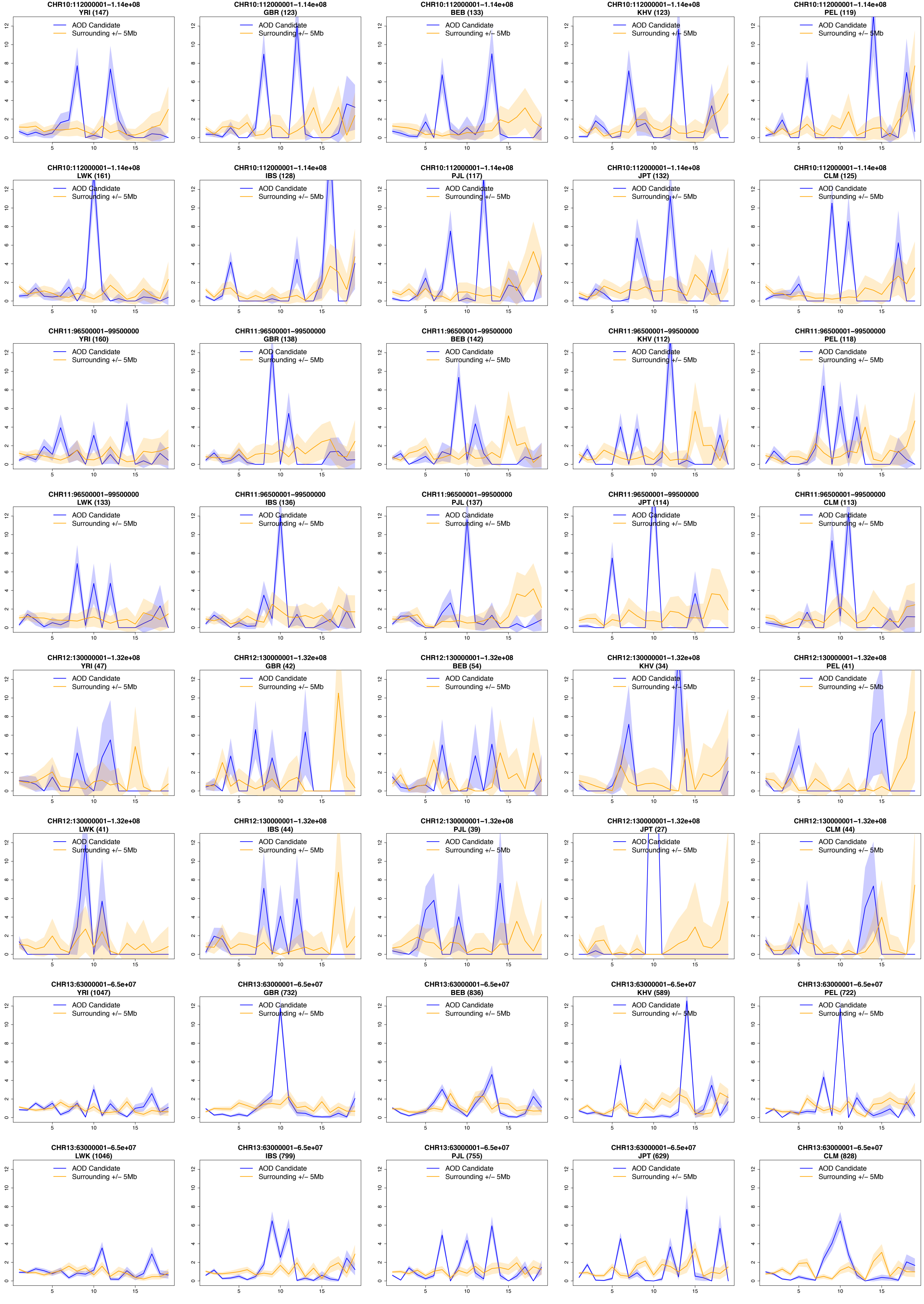
CHR10:112000001–1.14e+08 YRI (147)
CHR10:112000001–1.14e+08 GBR (123)
CHR10:112000001–1.14e+08 BEB (133)
CHR10:112000001–1.14e+08 KHV (123)
CHR10:112000001–1.14e+08 PEL (119)
CHR10:112000001–1.14e+08 LWK (161)
CHR10:112000001–1.14e+08 IBS (128)
CHR10:112000001–1.14e+08 PJL (117)
CHR10:112000001–1.14e+08 JPT (132)
CHR10:112000001–1.14e+08 CLM (125)
CHR11:96500001–99500000 YRI (160)
CHR11:96500001–99500000 GBR (138)
CHR11:96500001–99500000 BEB (142)
CHR11:96500001–99500000 KHV (112)
CHR11:96500001–99500000 PEL (118)
CHR11:96500001–99500000 LWK (133)
CHR11:96500001–99500000 IBS (136)
CHR11:96500001–99500000 PJL (137)
CHR11:96500001–99500000 JPT (114)
CHR11:96500001–99500000 CLM (113)
CHR12:130000001–1.32e+08 YRI (47)
CHR12:130000001–1.32e+08 GBR (42)
CHR12:130000001–1.32e+08 BEB (54)
CHR12:130000001–1.32e+08 KHV (34)
CHR12:130000001–1.32e+08 PEL (41)
CHR12:130000001–1.32e+08 LWK (41)
CHR12:130000001–1.32e+08 IBS (44)
CHR12:130000001–1.32e+08 PJL (39)
CHR12:130000001–1.32e+08 JPT (27)
CHR12:130000001–1.32e+08 CLM (44)
CHR13:63000001–6.5e+07 YRI (1047)
CHR13:63000001–6.5e+07 GBR (732)
CHR13:63000001–6.5e+07 BEB (836)
CHR13:63000001–6.5e+07 KHV (589)
CHR13:63000001–6.5e+07 PEL (722)
CHR13:63000001–6.5e+07 LWK (1046)
CHR13:63000001–6.5e+07 IBS (799)
CHR13:63000001–6.5e+07 PJL (755)
CHR13:63000001–6.5e+07 JPT (629)
CHR13:63000001–6.5e+07 CLM (828)
AOD Candidate
Surrounding +/– 5Mb

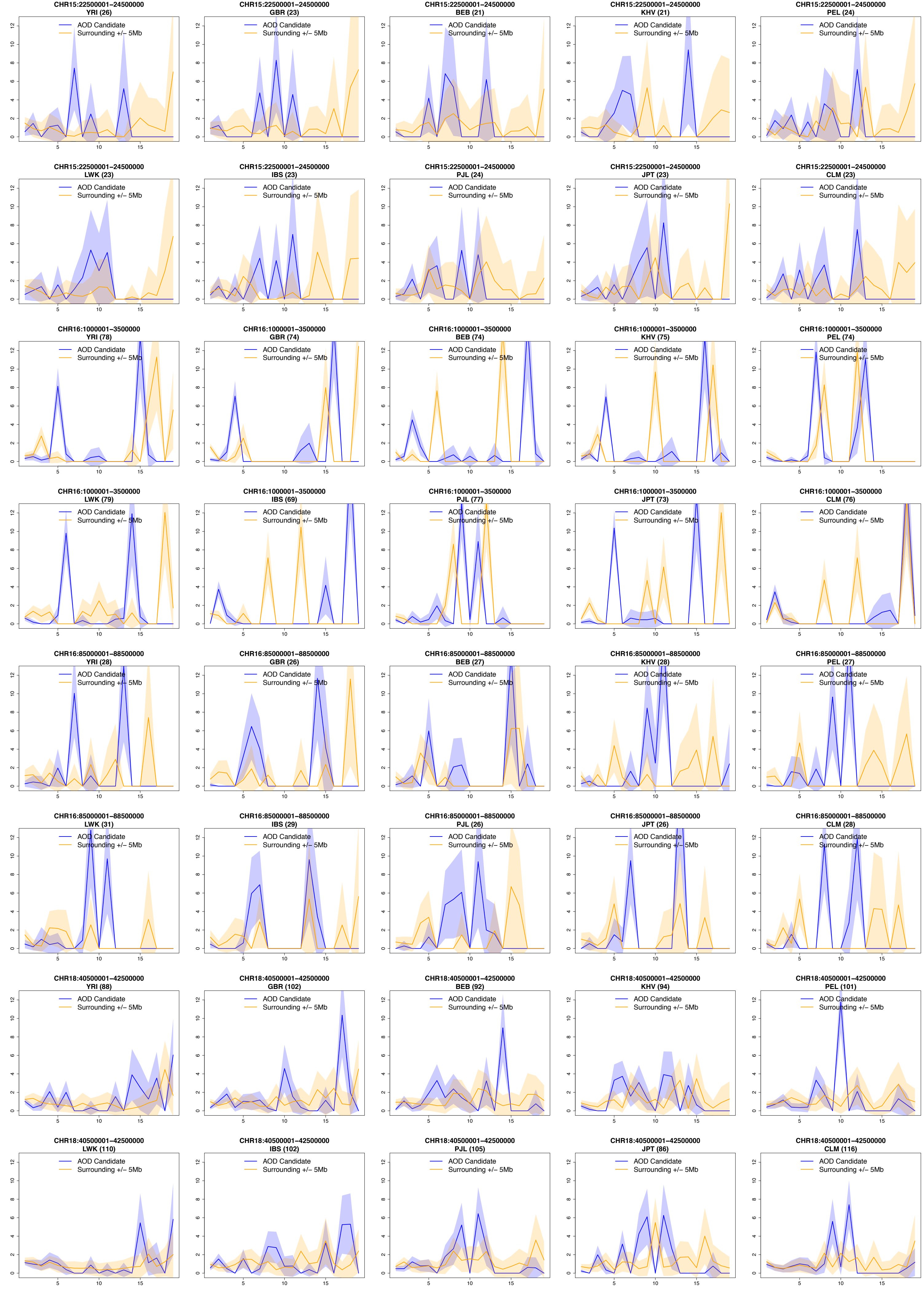
CHR15:22500001–24500000
YRI (26)
AOD Candidate
Surrounding +/– 5Mb
CHR15:22500001–24500000
GBR (23)
AOD Candidate
Surrounding +/– 5Mb
CHR15:22500001–24500000
BEB (21)
AOD Candidate
Surrounding +/– 5Mb
CHR15:22500001–24500000
KHV (21)
AOD Candidate
Surrounding +/– 5Mb
CHR15:22500001–24500000
PEL (24)
AOD Candidate
Surrounding +/– 5Mb
CHR15:22500001–24500000
LWK (23)
AOD Candidate
Surrounding +/– 5Mb
CHR15:22500001–24500000
IBS (23)
AOD Candidate
Surrounding +/– 5Mb
CHR15:22500001–24500000
PJL (24)
AOD Candidate
Surrounding +/– 5Mb
CHR15:22500001–24500000
JPT (23)
AOD Candidate
Surrounding +/– 5Mb
CHR15:22500001–24500000
CLM (23)
AOD Candidate
Surrounding +/– 5Mb
CHR16:1000001–3500000
YRI (78)
AOD Candidate
Surrounding +/– 5Mb
CHR16:1000001–3500000
GBR (74)
AOD Candidate
Surrounding +/– 5Mb
CHR16:1000001–3500000
BEB (74)
AOD Candidate
Surrounding +/– 5Mb
CHR16:1000001–3500000
KHV (75)
AOD Candidate
Surrounding +/– 5Mb
CHR16:1000001–3500000
PEL (74)
AOD Candidate
Surrounding +/– 5Mb
CHR16:1000001–3500000
LWK (79)
AOD Candidate
Surrounding +/– 5Mb
CHR16:1000001–3500000
IBS (69)
AOD Candidate
Surrounding +/– 5Mb
CHR16:1000001–3500000
PJL (77)
AOD Candidate
Surrounding +/– 5Mb
CHR16:1000001–3500000
JPT (73)
AOD Candidate
Surrounding +/– 5Mb
CHR16:1000001–3500000
CLM (76)
AOD Candidate
Surrounding +/– 5Mb
CHR16:85000001–88500000
YRI (28)
AOD Candidate
Surrounding +/– 5Mb
CHR16:85000001–88500000
GBR (26)
AOD Candidate
Surrounding +/– 5Mb
CHR16:85000001–88500000
BEB (27)
AOD Candidate
Surrounding +/– 5Mb
CHR16:85000001–88500000
KHV (28)
AOD Candidate
Surrounding +/– 5Mb
CHR16:85000001–88500000
PEL (27)
AOD Candidate
Surrounding +/– 5Mb
CHR16:85000001–88500000
LWK (31)
AOD Candidate
Surrounding +/– 5Mb
CHR16:85000001–88500000
IBS (29)
AOD Candidate
Surrounding +/– 5Mb
CHR16:85000001–88500000
PJL (26)
AOD Candidate
Surrounding +/– 5Mb
CHR16:85000001–88500000
JPT (26)
AOD Candidate
Surrounding +/– 5Mb
CHR16:85000001–88500000
CLM (28)
AOD Candidate
Surrounding +/– 5Mb
CHR18:40500001–42500000
YRI (88)
AOD Candidate
Surrounding +/– 5Mb
CHR18:40500001–42500000
GBR (102)
AOD Candidate
Surrounding +/– 5Mb
CHR18:40500001–42500000
BEB (92)
AOD Candidate
Surrounding +/– 5Mb
CHR18:40500001–42500000
KHV (94)
AOD Candidate
Surrounding +/– 5Mb
CHR18:40500001–42500000
PEL (101)
AOD Candidate
Surrounding +/– 5Mb
CHR18:40500001–42500000
LWK (110)
AOD Candidate
Surrounding +/– 5Mb
CHR18:40500001–42500000
IBS (102)
AOD Candidate
Surrounding +/– 5Mb
CHR18:40500001–42500000
PJL (105)
AOD Candidate
Surrounding +/– 5Mb
CHR18:40500001–42500000
JPT (86)
AOD Candidate
Surrounding +/– 5Mb
CHR18:40500001–42500000
CLM (116)
AOD Candidate
Surrounding +/– 5Mb

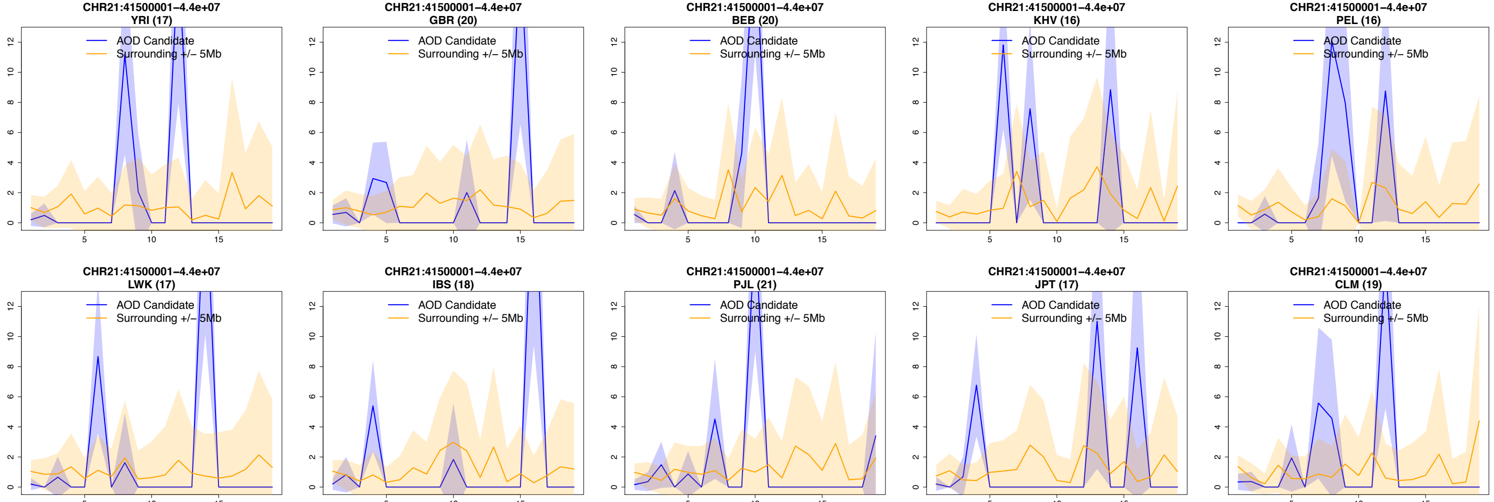
CHR21:41500001–4.4e+07
YRI (17)
AOD Candidate
Surrounding +/– 5Mb
CHR21:41500001–4.4e+07
GBR (20)
AOD Candidate
Surrounding +/– 5Mb
CHR21:41500001–4.4e+07
BEB (20)
AOD Candidate
Surrounding +/– 5Mb
CHR21:41500001–4.4e+07
KHV (16)
AOD Candidate
Surrounding +/– 5Mb
CHR21:41500001–4.4e+07
PEL (16)
AOD Candidate
Surrounding +/– 5Mb
CHR21:41500001–4.4e+07
LWK (17)
AOD Candidate
Surrounding +/– 5Mb
CHR21:41500001–4.4e+07
IBS (18)
AOD Candidate
Surrounding +/– 5Mb
CHR21:41500001–4.4e+07
PJL (21)
AOD Candidate
Surrounding +/– 5Mb
CHR21:41500001–4.4e+07
JPT (17)
AOD Candidate
Surrounding +/– 5Mb
CHR21:41500001–4.4e+07
CLM (19)
AOD Candidate
Surrounding +/– 5Mb

### 3 Item 2C : Heatmaps of the AOD candidate windows

Heatmaps for all 10 populations, 1 candidate region per page. Individuals are in rows, while low recombination variants are in columns (number of polymorphic SNPs change accross population). Homozygotes ancestral SNPs are in yellow, heterozygotes in orange and homozygotes derived in red. Clustering performed by heatmap2 in R, is more pronounced for non-Africans populations (no YRI nor LWK). The clustering has a similar pattern as a mixture of Fig. S4 panel A-C, even if we have only 10 individuals. Heatmaps are ordered the same way as in Item 2B.

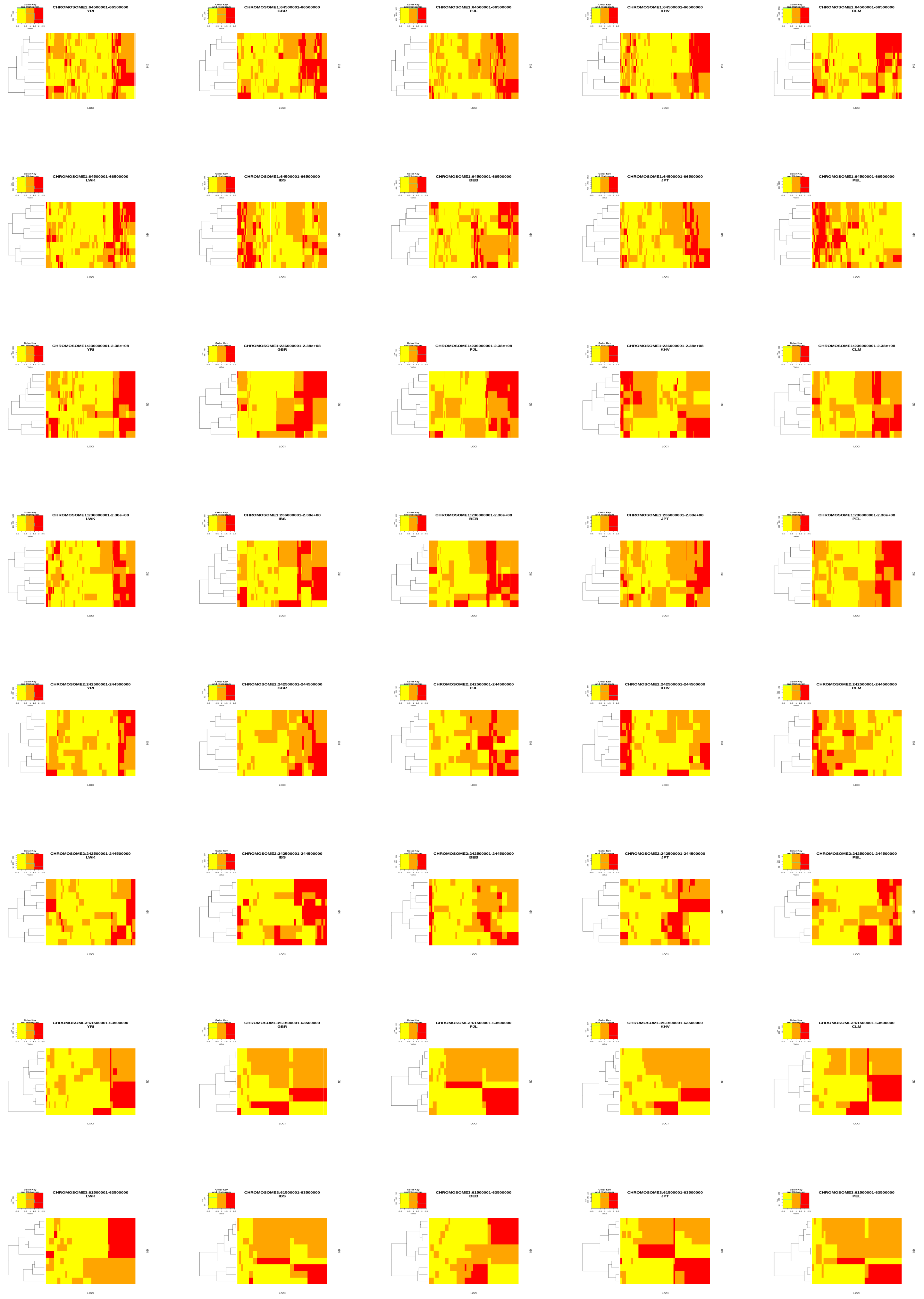
Color Key and Histogram
CHROMOSOME1:64500001-66500000 YRI
CHROMOSOME1:64500001-66500000 GBR
CHROMOSOME1:64500001-66500000 PJL
CHROMOSOME1:64500001-66500000 KHV
CHROMOSOME1:64500001-66500000 CLM
CHROMOSOME1:64500001-66500000 LWK
CHROMOSOME1:64500001-66500000 IBS
CHROMOSOME1:64500001-66500000 BEB
CHROMOSOME1:64500001-66500000 JPT
CHROMOSOME1:64500001-66500000 PEL
CHROMOSOME1:236000001-2.38e+08 YRI
CHROMOSOME1:236000001-2.38e+08 GBR
CHROMOSOME1:236000001-2.38e+08 PJL
CHROMOSOME1:236000001-2.38e+08 KHV
CHROMOSOME1:236000001-2.38e+08 CLM
CHROMOSOME1:236000001-2.38e+08 LWK
CHROMOSOME1:236000001-2.38e+08 IBS
CHROMOSOME1:236000001-2.38e+08 BEB
CHROMOSOME1:236000001-2.38e+08 JPT
CHROMOSOME1:236000001-2.38e+08 PEL
CHROMOSOME2:242500001-244500000 YRI
CHROMOSOME2:242500001-244500000 GBR
CHROMOSOME2:242500001-244500000 PJL
CHROMOSOME2:242500001-244500000 KHV
CHROMOSOME2:242500001-244500000 CLM
CHROMOSOME2:242500001-244500000 LWK
CHROMOSOME2:242500001-244500000 IBS
CHROMOSOME2:242500001-244500000 BEB
CHROMOSOME2:242500001-244500000 JPT
CHROMOSOME2:242500001-244500000 PEL
CHROMOSOME3:61500001-63500000 YRI
CHROMOSOME3:61500001-63500000 GBR
CHROMOSOME3:61500001-63500000 PJL
CHROMOSOME3:61500001-63500000 KHV
CHROMOSOME3:61500001-63500000 CLM
CHROMOSOME3:61500001-63500000 LWK
CHROMOSOME3:61500001-63500000 IBS
CHROMOSOME3:61500001-63500000 BEB
CHROMOSOME3:61500001-63500000 JPT
CHROMOSOME3:61500001-63500000 PEL
LOCI
IND

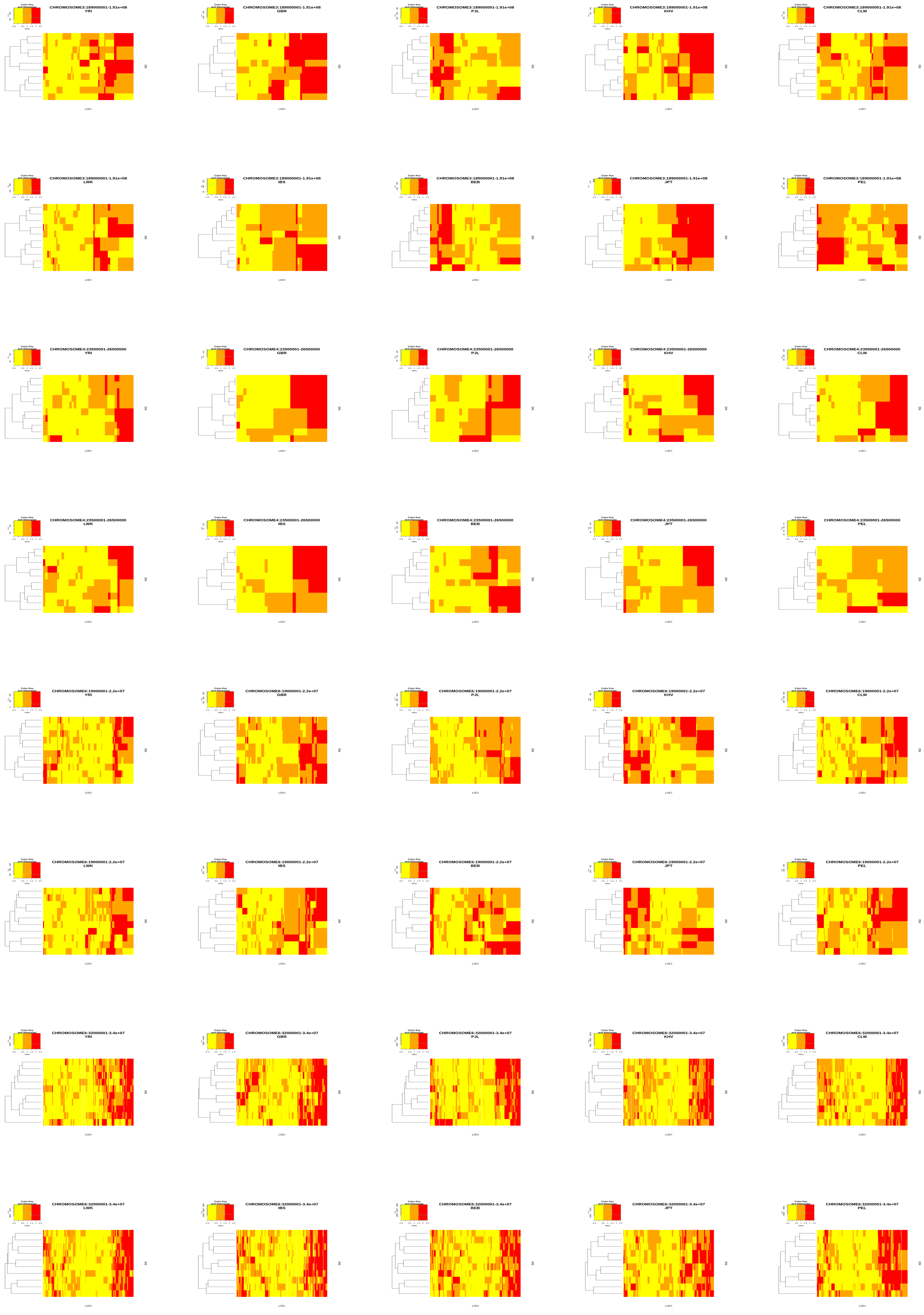
Color Key and Histogram
CHROMOSOME3:189000001-1.91e+08 YRI
CHROMOSOME3:189000001-1.91e+08 GBR
CHROMOSOME3:189000001-1.91e+08 PJL
CHROMOSOME3:189000001-1.91e+08 KHV
CHROMOSOME3:189000001-1.91e+08 CLM
CHROMOSOME3:189000001-1.91e+08 LWK
CHROMOSOME3:189000001-1.91e+08 IBS
CHROMOSOME3:189000001-1.91e+08 BEB
CHROMOSOME3:189000001-1.91e+08 JPT
CHROMOSOME3:189000001-1.91e+08 PEL
CHROMOSOME4:23500001-26500000 YRI
CHROMOSOME4:23500001-26500000 GBR
CHROMOSOME4:23500001-26500000 PJL
CHROMOSOME4:23500001-26500000 KHV
CHROMOSOME4:23500001-26500000 CLM
CHROMOSOME4:23500001-26500000 LWK
CHROMOSOME4:23500001-26500000 IBS
CHROMOSOME4:23500001-26500000 BEB
CHROMOSOME4:23500001-26500000 JPT
CHROMOSOME4:23500001-26500000 PEL
CHROMOSOME6:19000001-2.2e+07 YRI
CHROMOSOME6:19000001-2.2e+07 GBR
CHROMOSOME6:19000001-2.2e+07 PJL
CHROMOSOME6:19000001-2.2e+07 KHV
CHROMOSOME6:19000001-2.2e+07 CLM
CHROMOSOME6:19000001-2.2e+07 LWK
CHROMOSOME6:19000001-2.2e+07 IBS
CHROMOSOME6:19000001-2.2e+07 BEB
CHROMOSOME6:19000001-2.2e+07 JPT
CHROMOSOME6:19000001-2.2e+07 PEL
CHROMOSOME6:32000001-3.4e+07 YRI
CHROMOSOME6:32000001-3.4e+07 GBR
CHROMOSOME6:32000001-3.4e+07 PJL
CHROMOSOME6:32000001-3.4e+07 KHV
CHROMOSOME6:32000001-3.4e+07 CLM
CHROMOSOME6:32000001-3.4e+07 LWK
CHROMOSOME6:32000001-3.4e+07 IBS
CHROMOSOME6:32000001-3.4e+07 BEB
CHROMOSOME6:32000001-3.4e+07 JPT
CHROMOSOME6:32000001-3.4e+07 PEL
LOCI
ID

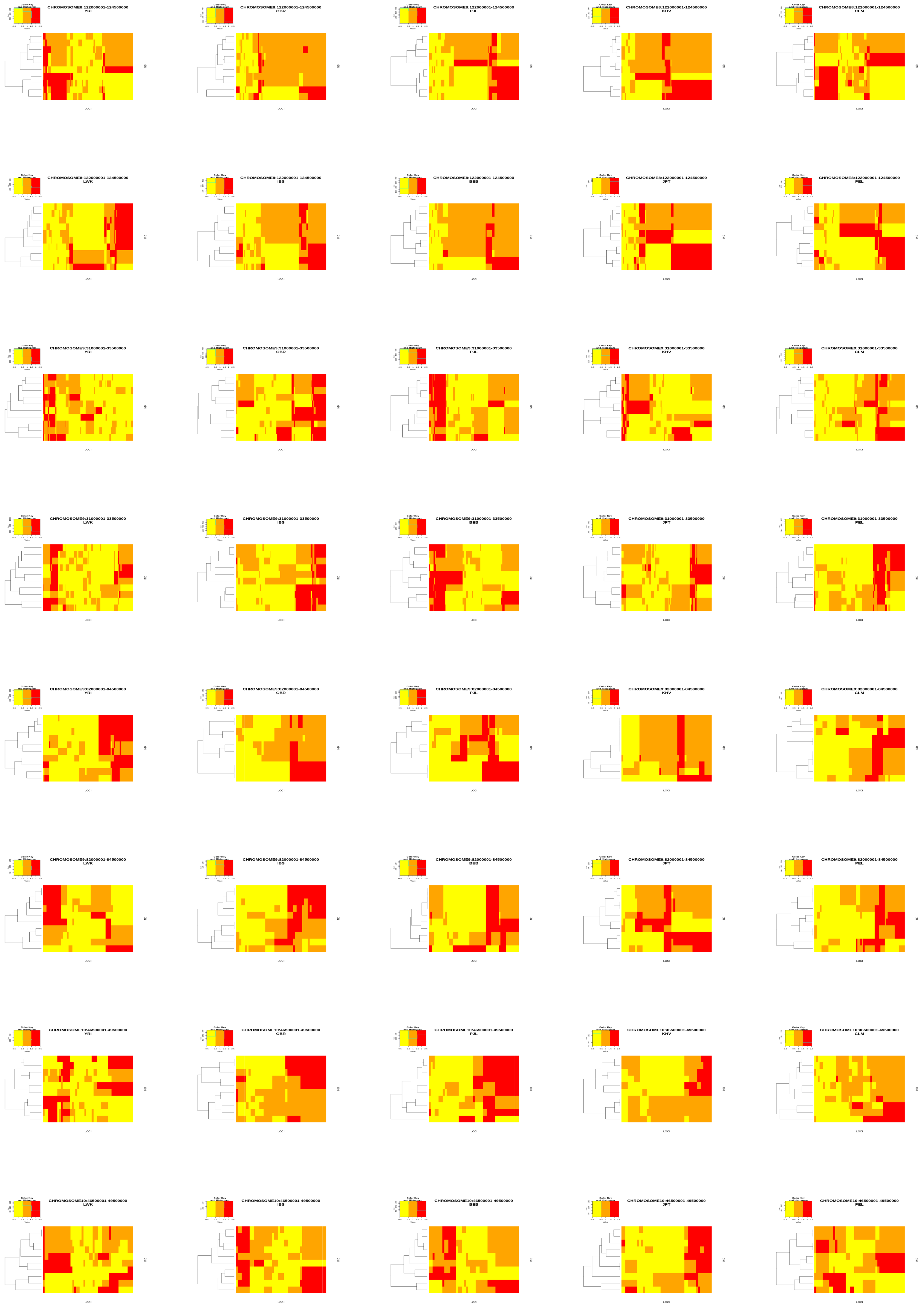
Color Key and Histogram
CHROMOSOME8:122000001-124500000 YRI
CHROMOSOME8:122000001-124500000 GBR
CHROMOSOME8:122000001-124500000 PJL
CHROMOSOME8:122000001-124500000 KHV
CHROMOSOME8:122000001-124500000 CLM
CHROMOSOME8:122000001-124500000 LWK
CHROMOSOME8:122000001-124500000 IBS
CHROMOSOME8:122000001-124500000 BEB
CHROMOSOME8:122000001-124500000 JPT
CHROMOSOME8:122000001-124500000 PEL
CHROMOSOME9:31000001-33500000 YRI
CHROMOSOME9:31000001-33500000 GBR
CHROMOSOME9:31000001-33500000 PJL
CHROMOSOME9:31000001-33500000 KHV
CHROMOSOME9:31000001-33500000 CLM
CHROMOSOME9:31000001-33500000 LWK
CHROMOSOME9:31000001-33500000 IBS
CHROMOSOME9:31000001-33500000 BEB
CHROMOSOME9:31000001-33500000 JPT
CHROMOSOME9:31000001-33500000 PEL
CHROMOSOME9:82000001-84500000 YRI
CHROMOSOME9:82000001-84500000 GBR
CHROMOSOME9:82000001-84500000 PJL
CHROMOSOME9:82000001-84500000 KHV
CHROMOSOME9:82000001-84500000 CLM
CHROMOSOME9:82000001-84500000 LWK
CHROMOSOME9:82000001-84500000 IBS
CHROMOSOME9:82000001-84500000 BEB
CHROMOSOME9:82000001-84500000 JPT
CHROMOSOME9:82000001-84500000 PEL
CHROMOSOME10:46500001-49500000 YRI
CHROMOSOME10:46500001-49500000 GBR
CHROMOSOME10:46500001-49500000 PJL
CHROMOSOME10:46500001-49500000 KHV
CHROMOSOME10:46500001-49500000 CLM
CHROMOSOME10:46500001-49500000 LWK
CHROMOSOME10:46500001-49500000 IBS
CHROMOSOME10:46500001-49500000 BEB
CHROMOSOME10:46500001-49500000 JPT
CHROMOSOME10:46500001-49500000 PEL
LOCI
IND

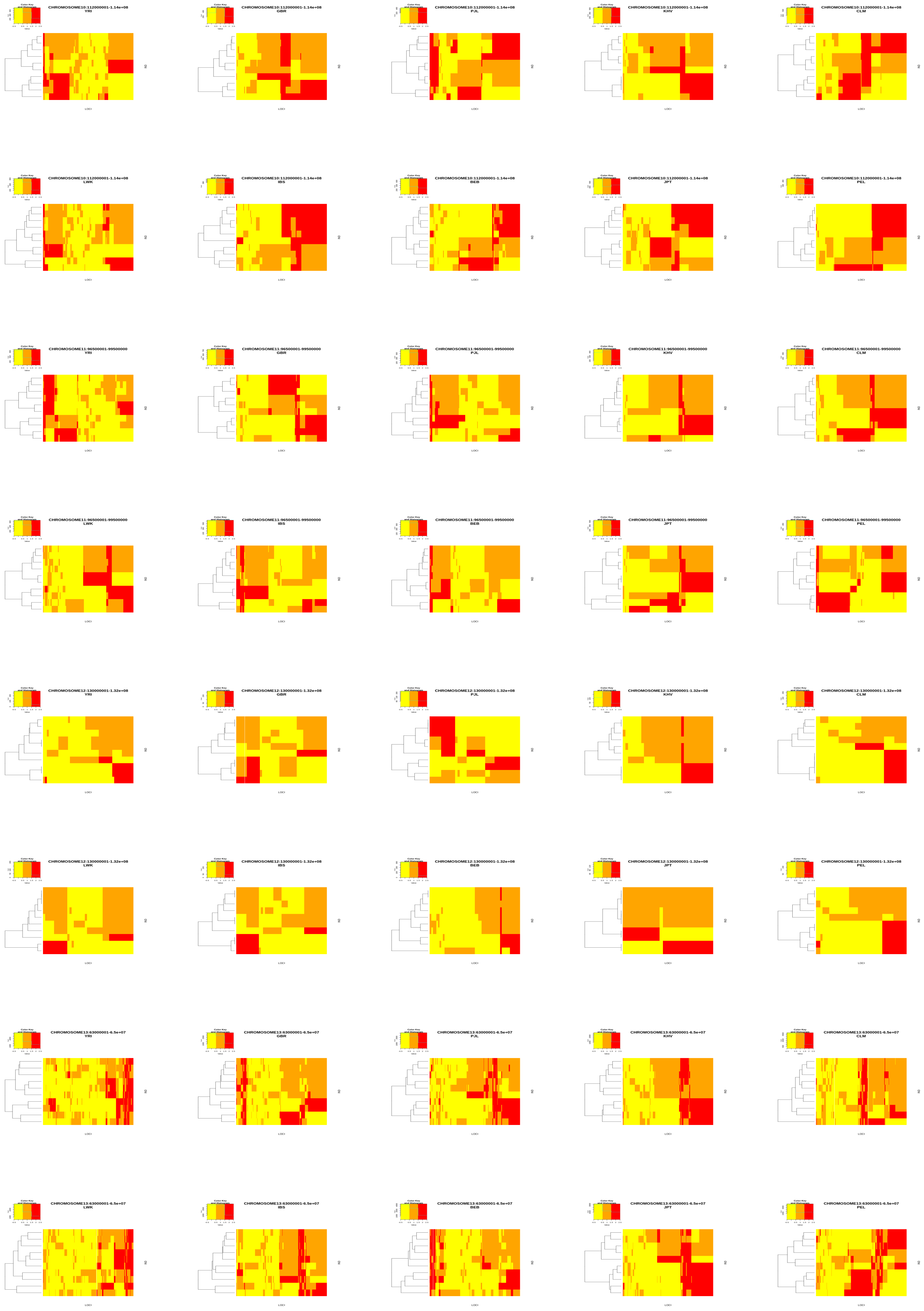
CHROMOSOME10:112000001-1.14e+08
YRI
CHROMOSOME10:112000001-1.14e+08
GBR
CHROMOSOME10:112000001-1.14e+08
PJL
CHROMOSOME10:112000001-1.14e+08
KHV
CHROMOSOME10:112000001-1.14e+08
CLM
CHROMOSOME10:112000001-1.14e+08
LWK
CHROMOSOME10:112000001-1.14e+08
IBS
CHROMOSOME10:112000001-1.14e+08
BEB
CHROMOSOME10:112000001-1.14e+08
JPT
CHROMOSOME10:112000001-1.14e+08
PEL
CHROMOSOME11:96500001-99500000
YRI
CHROMOSOME11:96500001-99500000
GBR
CHROMOSOME11:96500001-99500000
PJL
CHROMOSOME11:96500001-99500000
KHV
CHROMOSOME11:96500001-99500000
CLM
CHROMOSOME11:96500001-99500000
LWK
CHROMOSOME11:96500001-99500000
IBS
CHROMOSOME11:96500001-99500000
BEB
CHROMOSOME11:96500001-99500000
JPT
CHROMOSOME11:96500001-99500000
PEL
CHROMOSOME12:130000001-1.32e+08
YRI
CHROMOSOME12:130000001-1.32e+08
GBR
CHROMOSOME12:130000001-1.32e+08
PJL
CHROMOSOME12:130000001-1.32e+08
KHV
CHROMOSOME12:130000001-1.32e+08
CLM
CHROMOSOME12:130000001-1.32e+08
LWK
CHROMOSOME12:130000001-1.32e+08
IBS
CHROMOSOME12:130000001-1.32e+08
BEB
CHROMOSOME12:130000001-1.32e+08
JPT
CHROMOSOME12:130000001-1.32e+08
PEL
CHROMOSOME13:63000001-6.5e+07
YRI
CHROMOSOME13:63000001-6.5e+07
GBR
CHROMOSOME13:63000001-6.5e+07
PJL
CHROMOSOME13:63000001-6.5e+07
KHV
CHROMOSOME13:63000001-6.5e+07
CLM
CHROMOSOME13:63000001-6.5e+07
LWK
CHROMOSOME13:63000001-6.5e+07
IBS
CHROMOSOME13:63000001-6.5e+07
BEB
CHROMOSOME13:63000001-6.5e+07
JPT
CHROMOSOME13:63000001-6.5e+07
PEL
Color Key and Histogram
LOCI

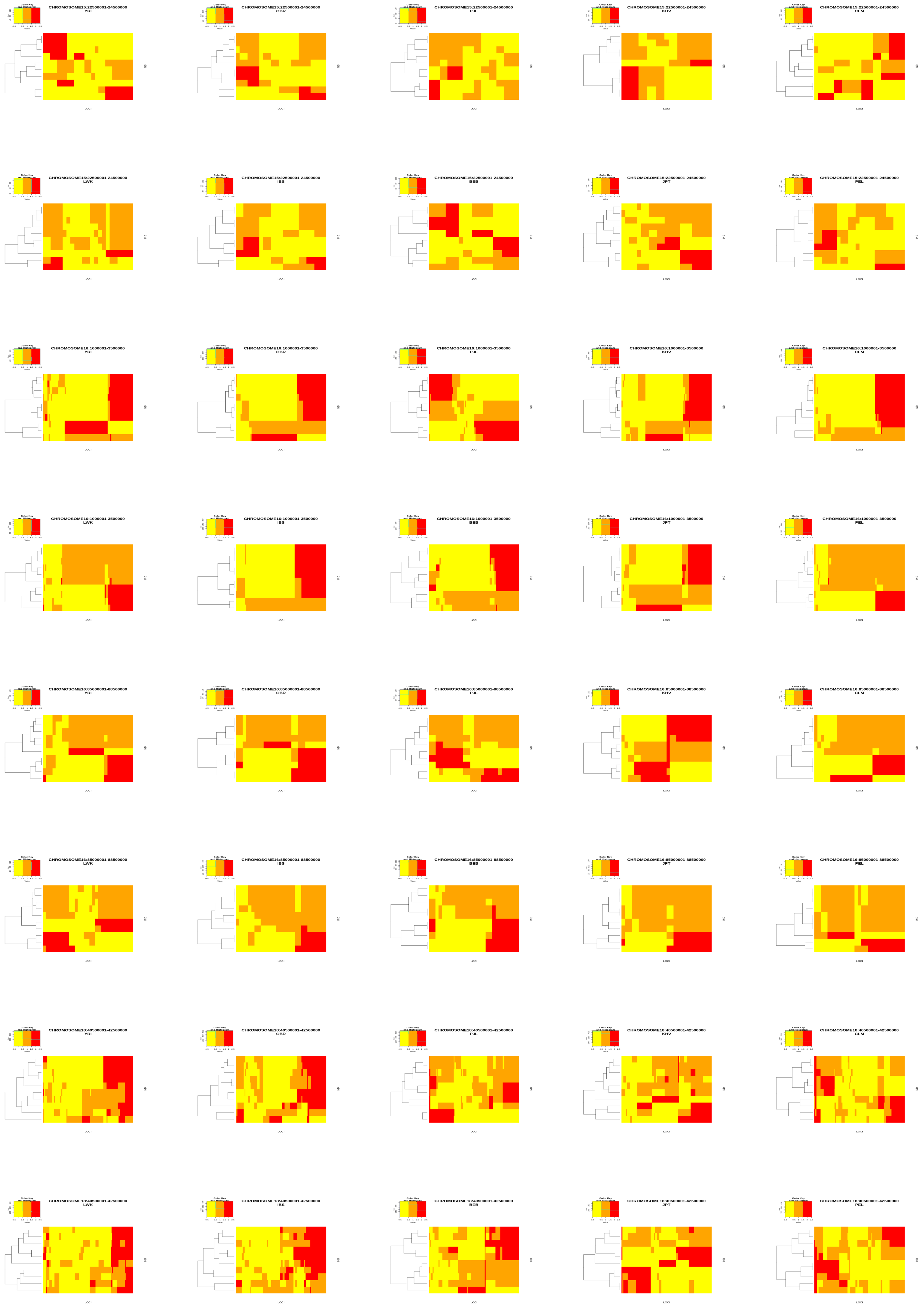

Color Key and Histogram
CHROMOSOME15:22500001-24500000 YRI
CHROMOSOME15:22500001-24500000 GBR
CHROMOSOME15:22500001-24500000 PJL
CHROMOSOME15:22500001-24500000 KHV
CHROMOSOME15:22500001-24500000 CLM
CHROMOSOME15:22500001-24500000 LWK
CHROMOSOME15:22500001-24500000 IBS
CHROMOSOME15:22500001-24500000 BEB
CHROMOSOME15:22500001-24500000 JPT
CHROMOSOME15:22500001-24500000 PEL
CHROMOSOME16:1000001-3500000 YRI
CHROMOSOME16:1000001-3500000 GBR
CHROMOSOME16:1000001-3500000 PJL
CHROMOSOME16:1000001-3500000 KHV
CHROMOSOME16:1000001-3500000 CLM
CHROMOSOME16:1000001-3500000 LWK
CHROMOSOME16:1000001-3500000 IBS
CHROMOSOME16:1000001-3500000 BEB
CHROMOSOME16:1000001-3500000 JPT
CHROMOSOME16:1000001-3500000 PEL
CHROMOSOME16:85000001-88500000 YRI
CHROMOSOME16:85000001-88500000 GBR
CHROMOSOME16:85000001-88500000 PJL
CHROMOSOME16:85000001-88500000 KHV
CHROMOSOME16:85000001-88500000 CLM
CHROMOSOME16:85000001-88500000 LWK
CHROMOSOME16:85000001-88500000 IBS
CHROMOSOME16:85000001-88500000 BEB
CHROMOSOME16:85000001-88500000 JPT
CHROMOSOME16:85000001-88500000 PEL
CHROMOSOME18:40500001-42500000 YRI
CHROMOSOME18:40500001-42500000 GBR
CHROMOSOME18:40500001-42500000 PJL
CHROMOSOME18:40500001-42500000 KHV
CHROMOSOME18:40500001-42500000 CLM
CHROMOSOME18:40500001-42500000 LWK
CHROMOSOME18:40500001-42500000 IBS
CHROMOSOME18:40500001-42500000 BEB
CHROMOSOME18:40500001-42500000 JPT
CHROMOSOME18:40500001-42500000 PEL
LOCI
ID

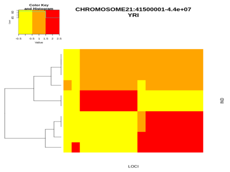
Color Key and Histogram
CHROMOSOME21:41500001-4.4e+07
YRI
LOCI

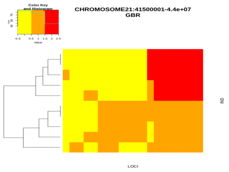
Color Key and Histogram
CHROMOSOME21:41500001-4.4e+07
GBR
LOCI

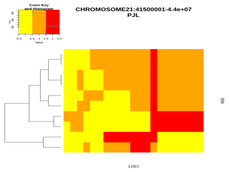
Color Key and Histogram
CHROMOSOME21:41500001-4.4e+07
PJL
LOCI

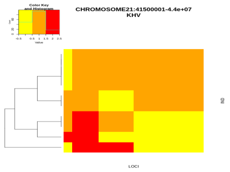
Color Key and Histogram
CHROMOSOME21:41500001-4.4e+07
KHV
LOCI

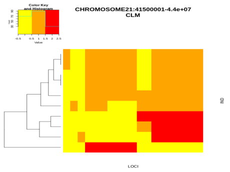
Color Key and Histogram
CHROMOSOME21:41500001-4.4e+07
CLM
LOCI

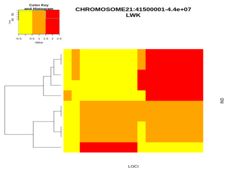
Color Key and Histogram
CHROMOSOME21:41500001-4.4e+07
LWK
LOCI

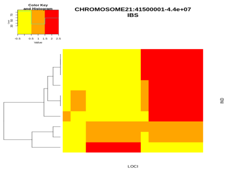
Color Key and Histogram
CHROMOSOME21:41500001-4.4e+07
IBS
LOCI

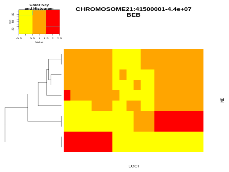
Color Key and Histogram
CHROMOSOME21:41500001-4.4e+07
BEB
LOCI

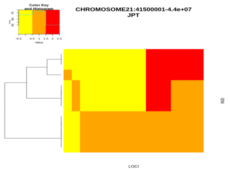
Color Key and Histogram
CHROMOSOME21:41500001-4.4e+07
JPT
LOCI

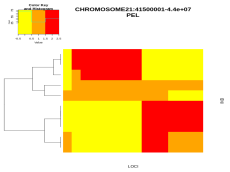
Color Key and Histogram
CHROMOSOME21:41500001-4.4e+07
PEL
LOCI
